# Loss of host factor-mediated m^6^Am methylation of the viral RNA cap impairs SARS CoV-2 replication

**DOI:** 10.64898/2026.04.10.717462

**Authors:** Radha Raman Pandey, Nadine Ebert, David Homolka, Tuba Barut, Bettina Salome Trüeb, Hanspeter Stalder, Elena Delfino, Cathrine Broberg Vågbø, Inês Berenguer Veiga, Sebastian Andreas Leidel, Volker Thiel, Ramesh S. Pillai

**Author notes:** These authors contributed equally (Radha Raman Pandey, Nadine Ebert). Correspondence: Ramesh S. Pillai, Volker Thiel.

## Abstract

Eukaryotic mRNAs are co-transcriptionally capped at the 5′ end with a methylated m^7^G moiety (cap0)^1^, which in higher eukaryotes is further methylated on the ribose (Nm) of the transcription start site (TSS) nucleotide to create the cap1 structure (m^7^GpppNm). Coronaviruses that replicate in the cytoplasm encode their own capping enzymes to acquire this cap1 structure which facilitates translation and shields them from the host innate immune system^2–5^. Here we report the identification of an additional *N^6^*-methyladenosine (m^6^A) methylation on the 5′ cap (m^7^Gpppm^6^Am) of the human coronavirus SARS-CoV-2 RNA. It is catalysed by the host m^6^A methylase PCIF1^6–9^ following capping by virus-encoded non-structural protein NSP 14 and NSP16^10^. Human cell cultures lacking *PCIF1* accumulate reduced levels of the viral RNA and support reduced viral replication. Furthermore, *Pcif1* mutant mice infected with SARS CoV-2 display milder symptoms. We identify the host RNA methyltransferase PCIF1 as a critical ally of SARS CoV-2 for viral replication.

---

The host nuclear CMTR1 installs the 2′-*O*-methyl mark (Nm) on the TSS nucleotide to generate the cap1 structure (m^7^GpppNm)^11–13^. When TSS nucleotide is an adenosine (Am), nuclear PCIF1 catalyses its *N^6^*-methylation to convert this into the di-methylated m^6^Am^6–9^. While the m^6^Am cap modification is dispensable for normal growth and fertility in mice^9^, it plays a role in stabilizing RNAs^8,9^ or regulating cap-dependent translation^6,7^. Host RNAs are also modified by the METTL3-METTL14 heterodimer^14–18^, which installs m^6^A marks internally at multiple locations. Such internal m^6^A marks can repel proteins that normally bind the unmethylated sequence, or are recognized by reader proteins, to regulate transcript splicing, stability and translation^19–23^.

Viruses that replicate in the cytoplasm, like the West Nile Virus and Coronaviruses, encode their own capping enzymes to produce RNAs with the cap1 structure^10^. Virus mutants that lack the 2′-*O*-methyltransferase trigger the innate immune pathway, as the cap0 viral RNAs are recognized by cytosolic sensors like RIG-I^2,3^ or MDA5^5^, triggering the interferon (IFN) production. This in turn leads to expression of IFN-stimulated genes (ISGs) that supress translation of the cap0 RNAs, causing viral attenuation^5,24,25^. Thus, virus-encoded cap1 is an essential survival mechanism for cytosolic viruses.

Viruses also hijack host factors for acquiring RNA modifications. The human metapneumovirus (MHPV)^26^, the Vesicular Stomatitis Virus (VSV)^27^, and many more, acquire internal m^6^A marks deposited by host METTL3. This helps to avoid detection by the sensor protein RIG-I, thereby reducing interferon expression. Likewise, the severe acute respiratory syndrome coronavirus 2 (SARS CoV-2) that is responsible for the current coronavirus disease 2019 (COVID-19) pandemic is also shown to be internally methylated by METTL3 at several sites^28–31^. These modifications are shown to help avoid detection by RIG-I^31^, promoting SARS CoV-2 replication^28,29^.

We wished to examine if there are additional RNA modifications on the SARS CoV-2 RNA. Coronaviruses like SARS CoV-2 have a positive-sense m^7^G-capped and polyadenylated ∼30 kb RNA as their genome^10^. Once the virus enters the host cell this genome is released into the cytoplasm where it behaves like a giant messenger RNA encoding for the two large overlapping ORFs (1a and 1b) encoding for non-structural proteins (NSP) 1-16 necessary for viral transcription and replication. Also produced are a further series of sub-genomic RNAs that are required for translation of the structural and accessory protein ORFs present at the C-term of the genome. To obtain viral RNA, we used African green monkey kidney-derived Vero E6 cells infected with a recombinant preparation of SARS CoV-2^32^ that is identical in sequence to the index SARS-CoV-2 (Wuhan-Hu-1).

Total RNA was isolated from infected cells and viral RNA was purified using a two-step protocol (Fig 1a). Purification of polyA+ RNA and deep sequencing shows that viral RNA constitutes ∼70% of this population (Fig. 1b). Further affinity-purification using virus-specific antisense capture probes (Supplementary Table 1) enriches viral RNAs to ∼90% of this preparation (Fig. 1b). We used this highly enriched fraction for mass spectrometry (Fig. 1c) and detected RNA modifications separately for the body of the RNA (internal only), and the entire RNA (cap+ internal) (Fig. 1d). Demonstrating the specificity of this protocol, m^7^G is detected only in the complete RNA fraction (cap+ internal) (Fig. 1e). Likewise, an increased abundance of ribose-methylated adenosine (Am) is observed in the complete RNA (cap+ internal) fraction (Fig. 1e and Extended Data Fig. 1a). This is consistent with the cap1 structure and the presence of an adenosine as the 5’-terminal nucleotide of SARS CoV-2 RNA. We also detect m^6^A and the dimethylated m^6^Am (*N^6^*-methyl-2′-*O*-methyl adenosine) (Fig. 1e). Interestingly, m^6^Am is detected only in the complete (cap+ internal) fraction, and not on the body of the RNA (internal only), suggesting its presence on the SARS CoV-2 viral RNA cap structure. Note that the purified viral RNA has higher levels of m^6^Am, compared to the polyA+ fraction which still has some host mRNAs (Fig. 1b and 1e). Four independent studies recently mapped the presence of internal m^6^A modification at several positions along the length of the SARS CoV-2 RNA^28–31^, but here we identify m^6^Am as an additional viral cap modification that was not previously reported.

**Fig. 1.**
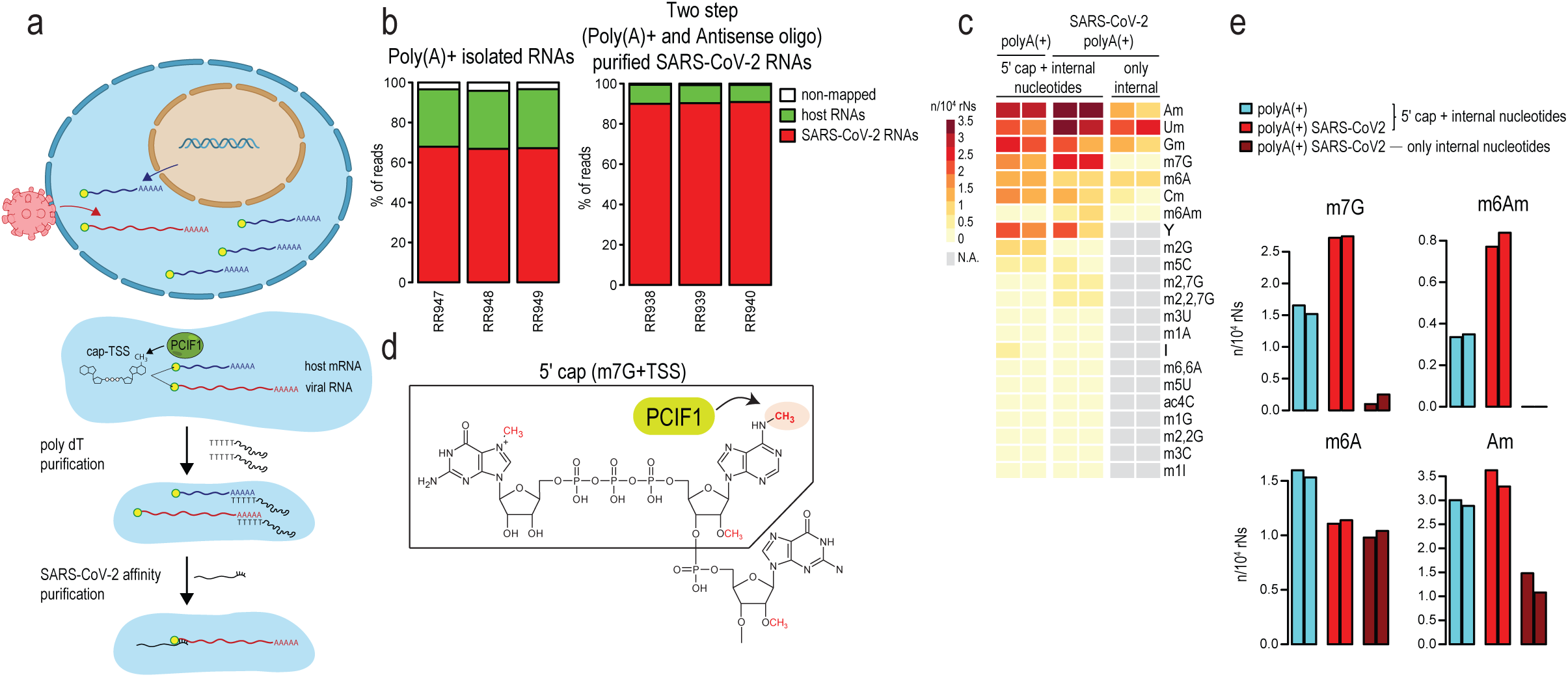
RNA modifications of SARS CoV-2. **a**, Two-step protocol for purification of SARS CoV-2 RNA from infected Vero E6 cells by polyA+ and subsequent antisense oligo affinity enrichment. **b**, SARS CoV-2 RNA constitutes 70% of the polyA+ fraction and 90% of the antisense oligo-enriched fraction. Experiments done in triplicates. **c**, Heatmap showing enrichment of RNA modifications in the different purified fractions. Modifications on the entire RNA (cap+ internal) or within the body of the RNA minus m^7^G+TSS (internal only) are indicated separately. Experiments done in duplicates. **d**, Chemical structure of the 5′ cap found on eukaryotic RNA pol II transcripts. The m^7^G moiety is attached via a triphosphate bridge to the transcription start site (TSS) nucleotide (cap0). Ribose methylation of the TSS adenosine (cap1), and its *N^6^* methylation by PCIF1 to form m^6^Am are shown. The boxed region indicates the 5′ cap (m^7^G+TSS) that is absent in the “internal only” fraction analyzed by RNA mass spectrometry in panel c. **e**, Bar plots show abundance (number of modified nucleotides/10^4^ nucleotides) of some of the modifications. Notice the absence of m^7^G and m^6^Am in the “internal only” fraction, indicating that they are exclusively part of the 5′ cap (m^7^G+TSS) structure. Experiments done in duplicates.

To map the m^6^A sites on the SARS CoV-2 RNA, we used m^6^A-IPseq that depends on the use of anti-m^6^A antibodies that do not distinguish between m^6^A and m^6^Am. PolyA+ RNA isolated from infected Vero E6 cells were fragmented and subjected to immunoprecipitation with anti-m^6^A antibodies. Mapping of the reads to the viral genome and calculation of enrichment after immunoprecipitation (m^6^A reads/input) reveals a strong peak at the 5′ end of the viral genome (Fig. 2a and Extended Data Fig. 2a). Additional smaller peaks were found along the length of the RNA, but more towards the 3′ end. Although the protocol suffers from lack of nucleotide-resolution, based on the reads with their 5′ end on first nucleotide having the highest enrichment, we propose that the 5′ end peak corresponds largely to the m^6^Am modification of the 5′-terminal nucleotide. A similar m^6^A-IPseq analysis with RNA isolated from human hepatoma cell line Huh7 infected with the common cold-causing human coronavirus hCoV-229E also reveals an identical peak at the 5′ end (Fig. 2b and Extended Data Fig. 2b). Both SARS CoV-2 and hCoV-229E have an adenosine as the 5’-terminal nucleotide. These results are consistent with the presence of a cap-specific m^6^Am modification on SARS CoV-2 and hCoV-229E RNAs.

**Fig. 2.**
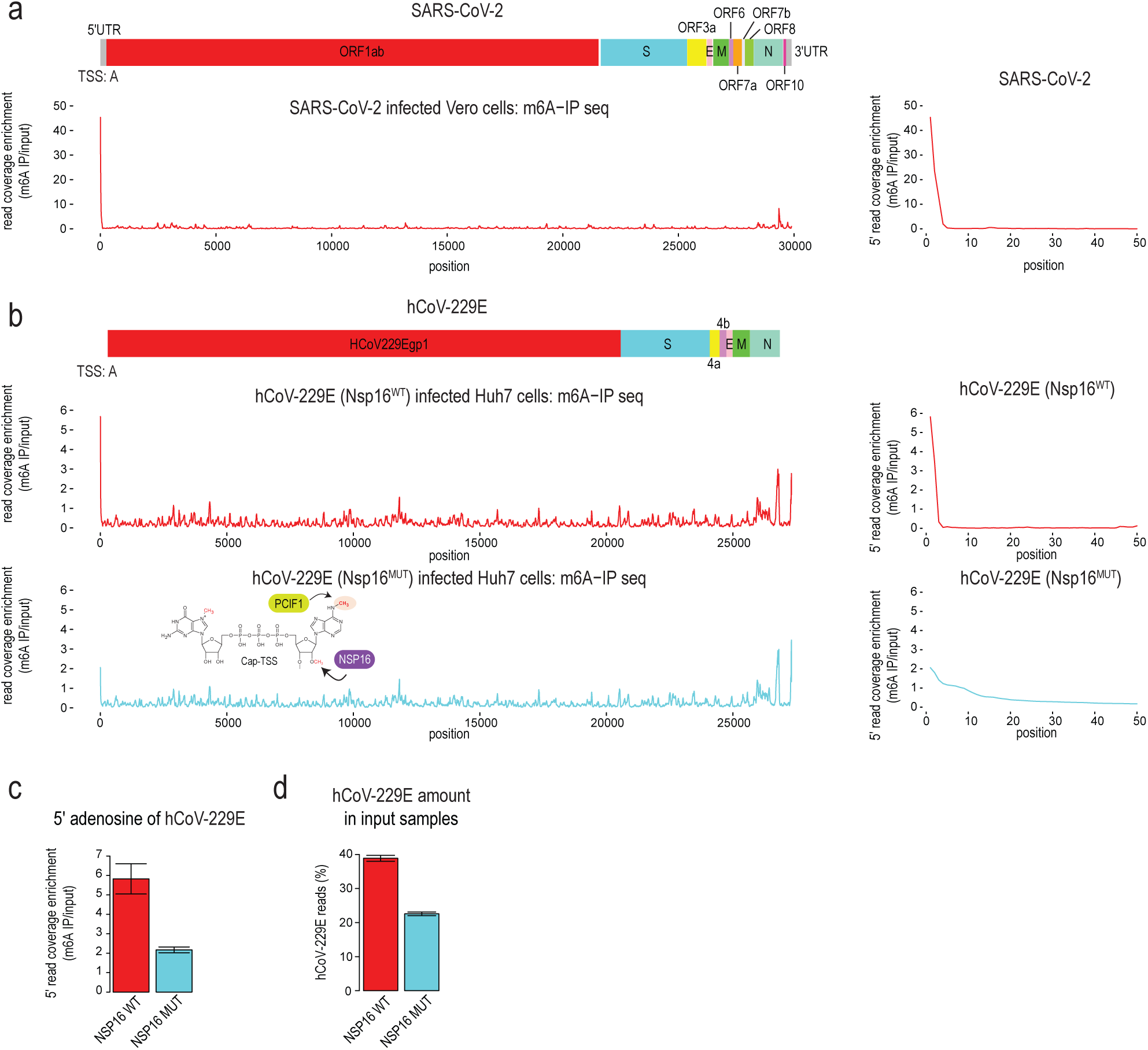
m^6^Am on coronavirus RNA cap and its dependence on viral NSP16. **a**, Mapping of m^6^A on SARS CoV-2 genome using m^6^A-IPseq with RNA from infected Vero E6 cells. The viral genome organization is shown on the top. The averaged read coverage enrichments in m^6^A-IPseq libraries from three replicas are plotted along the genome. The prominent 5′ end peak corresponds to m^6^Am as the coronavirus has a cap1 structure (Am). The 5′ end read enrichment is shown on the right only for the 5′ end of the viral genome. **b**, Mapping of m^6^A on the common cold-causing human coronavirus hCoV-229E using m^6^A-IP seq (three replicas) with RNA from infected Huh7 cells. The prominent 5′ end peak corresponding to m^6^Am is reduced when cells are infected with a virus mutant for the virus-encoded NSP16 that catalyzes 2′-*O*-methylation of the TSS nucleotide (Am) (three replicas). **c**, Bar plot showing enrichment in m^6^A-IPseq libraries for the reads starting at the 5′ adenosine, which demonstrates a reduction in m^6^Am in the *NSP16* mutant hCoV-229E. Data are presented as the mean ± s.d. (n = 3 replicas) **d**, Bar plot showing proportion of reads mapping to the viral genome which demonstrates the reduced viral RNA levels when cells are infected with the NSP16 mutant. Data are presented as the mean ± s.d. (n = 3 replicas).

Biochemical studies have shown that a complex consisting of the SARS coronavirus non-structural protein16 (NSP16) and its activator protein NSP10 is responsible for 2′-*O*-methylation of the 5′-terminal adenosine into Am to create the cap1 structure^33^, with NSP16 being the catalytically active component^34^. This Am moiety may represent the building block for further *N^6^*-methylation to produce the m^6^Am mark. To test its requirement for the m^6^A methylation, we used a *NSP16* mutant hCoV-229E^5^. Mapping of m^6^A-IPseq reads on RNA isolated from Huh7 cells infected with the mutant virus (Fig. 2b and Extended Data Fig. 2b) reveals a ∼3-fold reduction in the 5′ end peak (Fig. 2c and Extended Data Fig. 2c), indicating that prior ribose methylation (Am) of the 5′-terminal adenosine is a pre-requisite for efficient formation of m^6^Am on the cap structure. Consistent with previous reports of viral attenuation due to loss of *NSP16*^5,24^, the mutant hCoV-229E accumulates ∼50% less RNA overall when compared to the wildtype virus (Fig. 2d and Extended Data Fig. 2d). Taken together, these results show that NSP16-mediated cap1 formation is required for robust m^6^Am methylation of the coronavirus cap structure.

Nuclear PCIF1 is the cap-specific mammalian RNA methyltransferase that catalyse *N^6^*-methylation of the TSS adenosine of host RNA pol II transcripts^6^. To investigate its involvement in viral RNA methylation, we used wildtype and *PCIF1* knockout (KO) human HEK293T cells that were made suitable for infection with SARS CoV-2 by stably expressing the human Angiotensin-converting enzyme 2 (ACE2), the host cell surface receptor required for virus attachment and entry^35^. The cell lines were infected with SARS CoV-2 and 48h post-infection, polyA+ RNA was extracted and subject to m^6^A-IPseq (Fig. 3a). Consistent with a role for PCIF1 in cap methylation of the viral RNA, a ∼3-fold reduction in the 5′ end m^6^A peak is observed in the *PCIF1* KO cells (Fig. 3b and Extended Data Fig. 3a). Mass spectrometry analysis of polyA+ RNA extracted from the cells show a complete absence of m^6^Am methylation in the *PCIF1* KO cells (Fig. 3c and Extended Data Fig. 3b). Given this, the residual peak at the 5′ end in the *PCIF1* KO cells is likely due to other internal m^6^A modifications present closer to the 5′ end.

**Fig. 3.**
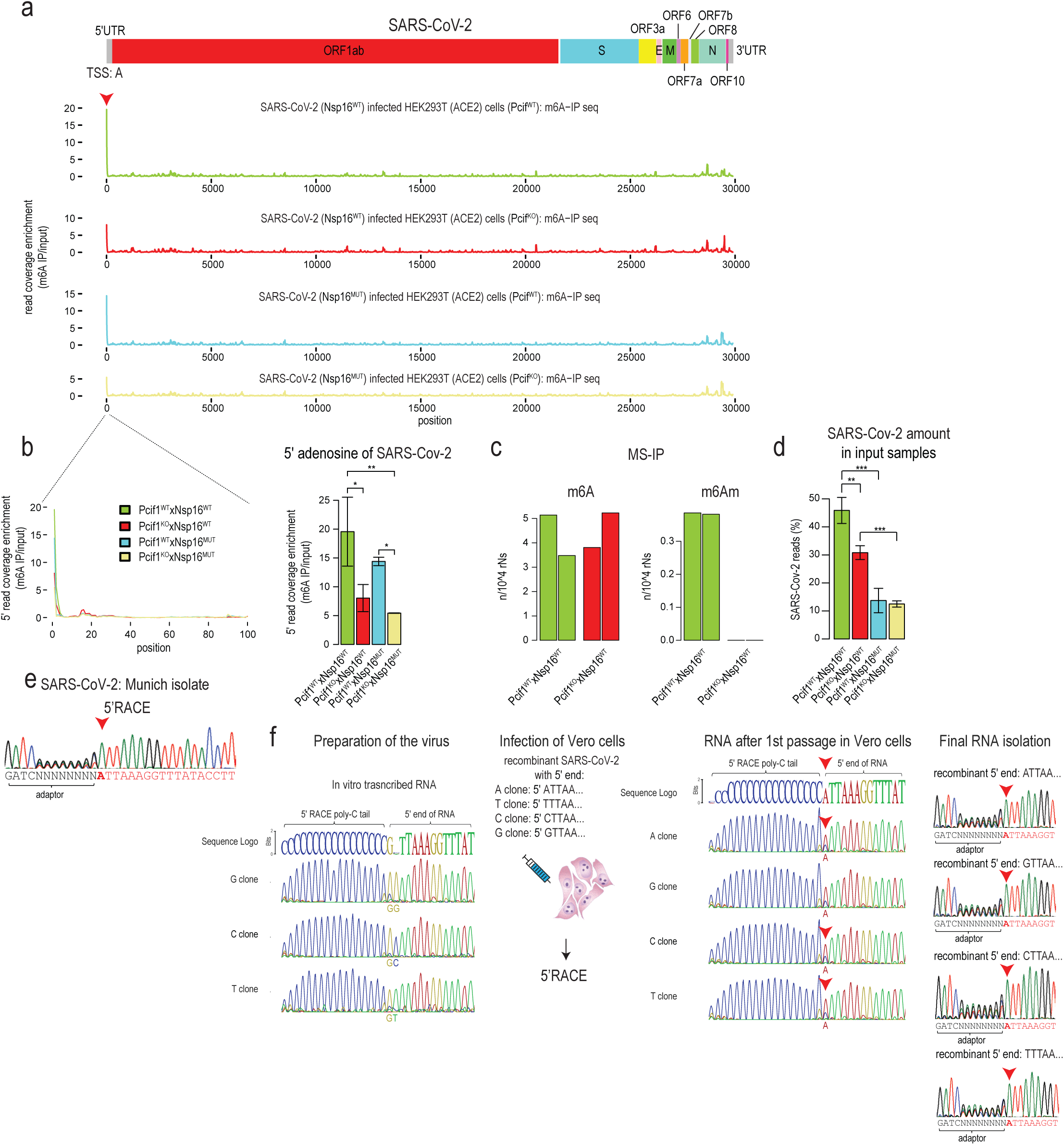
Immutability of the 5′-terminal nucleotide of SARS CoV-2 and its m^6^Am methylation by host PCIF1. **a**, Mapping of m^6^A on SARS CoV-2 genome using m^6^A-IPseq with RNA from infected wildtype or *PCIF1* KO human HEK293T ACE2 cells. Viruses used were either wildtype SARS CoV-2 or a mutant lacking *NSP16*. The viral genome organization is shown on the top. The averaged read coverage enrichments in m^6^A-IPseq libraries from three replicas are plotted along the genome. The prominent 5′ end peak corresponds to m^6^Am as the coronavirus has a cap1 structure (Am). **b**, The 5′ end read enrichment is shown only for the 5′ end of the viral genome. Bar plot shows enrichment in m^6^A-IPseq libraries for the reads starting at the 5′ adenosine. A reduction is seen in the *PCIF1* KO HEK293T ACE2 cells. Data are presented as the mean ± s.d. (n = 3 replicas); The p values obtained by Tukey’s HSD after ANOVA. ∗p ≤ 0.05, ∗∗p ≤ 0.01, ∗∗∗p ≤ 0.001. **c**, RNA mass spectrometry of polyA+ RNA from wildtype or *PCIF1* KO HEK293T ACE2 cells showing the absence of m^6^Am in the KO cells. Experiments done in duplicates. **d**, Bar plot showing proportion of reads mapping to the viral genome which demonstrates reduced viral RNA levels in the *PCIF1* KO HEK293T ACE2 cells. There is also the expected decrease in RNA levels in the *NSP16* mutant virus. Data are presented as the mean ± s.d. (n = 3 replicas); The p values obtained by Tukey’s HSD after ANOVA. ∗p ≤ 0.05, ∗∗p ≤ 0.01, ∗∗∗p ≤ 0.001 **e**, 5′ RACE analysis of 5′-terminal nucleotide of a natural isolate of SARS CoV-2. An adenosine is the 5′-terminal nucleotide. **f**, 5′ RACE analysis of the in vitro transcribed RNA templates with different viral start sites nucleotides, and that of the recombinant SARS CoV-2 viruses produced from these. All mutant templates result in production of revertants that have an adenosine as the viral 5′-terminal nucleotide (red arrowhead).

In vitro experiments reveal that a prior 2′-*O*-methylation (Am) of the substrate is not essential for PCIF1 activity, but can enhance the kinetics of methylation^6-8^. Consistently, mutation of NSP16 in the SARS CoV-2 genome results in only a small reduction (∼1.5-fold) in the 5′ end peak in wildtype cells (Fig. 3b). Thus, m^6^Am methylation by PCIF1 on SARS CoV-2 cap structure may not be dependent on prior ribose methylation by virus-encoded NSP16, but its activity may present an optimal substrate for PCIF1 function. Quantifying the overall viral RNA levels in these studies provides support for two main conclusions (Fig. 3d and Extended Fig. 3c). First, as expected for coronaviruses^5,24^, mutation of virus-encoded NSP16 results in an attenuated state with ∼4-fold reduction in SARS CoV-2 RNA. Analysis of host gene expression in the infected cells did not reveal an activation of the interferon pathway, but GO term analysis implicates an alteration in genes involved in host transcription and translation (Extended Fig. 3d-f). Second, absence of host PCIF1 also reduces viral RNA levels, albeit at a lower level (∼1.5-fold). Taken together, the SARS CoV-2 cap1 structure synthesized by viral enzymes is further modified by host PCIF1 to install an m^6^Am modification, and this permits robust accumulation of the viral RNA.

The recent pandemic has seen the SARS CoV-2 mutate at multiple internal positions generating several variants of concern as defined by the World Health Organization (WHO): alpha, beta, gamma, delta, and the latest omicron. Given the importance of the 5′-terminal adenosine for PCIF-1-mediated catalysis of m^6^Am, we asked whether SARS CoV-2 replication can be sustained when 5′-terminal nucleotide is mutated. This is an important question given that several other coronaviruses like the mouse hepatitis virus (MHV) use a guanosine as the 5’-terminal nucleotide^36^.

A clinical isolate of SARS CoV-2 has an adenosine as the 5′-terminal nucleotide as detected by 5′ RACE experiments (Fig. 3e). We prepared T7-transcribed 30 kb viral RNA preparations where the 5′-terminal nucleotide is mutated to a guanosine, cytidine or uridine (Fig. 3f). These were used for virus production after electroporation into mammalian cell cultures, as previously described^32^. As expected, 5′ RACE showed the correct replacement of the 5′-terminal nucleotide in all in vitro transcribed constructs (Fig. 3f). Within the initial culture, we observed the expected cytopathic effect (CPE) after 2-3 days only for the wild-type construct possessing the authentic 5′-terminal adenosine. This is consistent with robust virus production. Notably, constructs containing 5′-terminal guanosine, cytidine, or uridine, gave rise to a delayed CPE appearance after 5-7 days. Supernatants from these initial cultures were transferred to fresh VeroE6 cells to prepare RNA for assessment of the 5′-terminal sequences. Surprisingly, 5′ RACE experiments show that all the templates result in the rescued viruses being revertants that start with an adenosine (m^7^Gppp**<u>A</u>**TTAAA-) (Fig. 3f). This shows that some specific requirement pressures SARS CoV-2 to maintain a 5′-terminal adenosine for efficient replication. Inability of NPS16/NSP10 to efficiently methylate a 5′-terminal other then adenosine^37^ together with lack of 2′-*O* methylation (cap0) leading to attenuation^2-5^, might explain the selection of revertants with a 5′-terminal adenosine. Our identification of cap-specific m^6^Am and its role in boosting viral RNA levels in vivo may be an additional reason for immutability of the 5′-terminal adenosine of SARS CoV-2.

To examine the impact of host PCIF1 on viral replication, we used A549 human alveolar basal epithelial cells with reduced PCIF1 expression (Fig. 4a). The cells were made competent for SARS CoV-2 infection by stable expression of the hACE2 receptor. These cells were chosen as they facilitated a much more robust viral replication compared to the HEK293 ACE2 cells. Western analysis shows reduced expression of PCIF1 in the mutant cell line (Fig. 4a). We infected either A459 hACE2 WT or *PCIF1* mut cells with 10^4^ PFU/ml of the virus and monitored viral titres daily, over a period of three days. In triplicate, independent experiments, we noticed a dramatic decrease by orders of magnitude (wildtype= 10^2^ PFU/ml vs mutant=10^3^ to 10^4^ PFU/ml) in viral titres in the *PCIF1* mutant cells (Fig. 4a and Extended Data Fig. 4a). Taken together with the fact that viral caps are decorated with m^6^Am methylation, and that this is important for robust viral RNA accumulation, these results place PCIF1 as a critical host factor that can contribute to efficient viral replication.

**Fig. 4.**
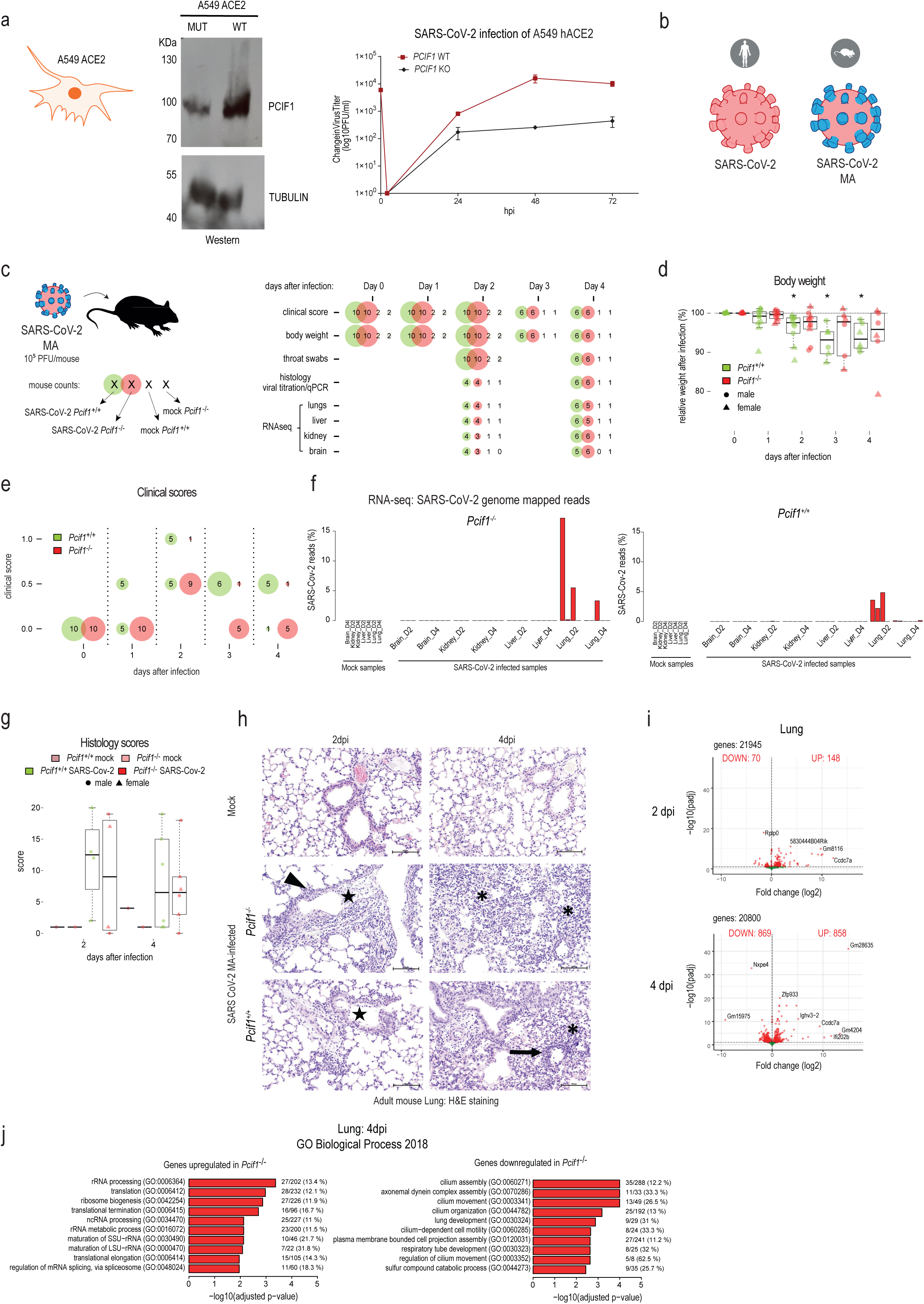
SARS CoV-2 replication is reduced in the absence of host PCIF1. **a**, Infection of A459 ACE2 WT and PCIF1-depleted (Mut) cells with SARS CoV-2. Western blot analysis of PCIF1 in the WT and Mut cells. Viral titres estimated by plaque assays. Data are presented as the mean ± s.d. (n = 3 replicas). **b**, A mouse-adapted version of human SARS CoV-2 coronavirus with mutations (SARS CoV-2 MA10) in the spike that allows attachment to the mouse ACE2 receptor was used for mouse infection studies. **c**, Either wildtype (n=10) or *Pcif1* KO (n=10) mice were infected with SARS CoV-2 MA at day 0 and followed for a period of 4 days. Mock-treated animals were left uninfected. **d**, Body weight was measured daily. The p-values were obtained using one-tailed one-sample t.test of mean relative body weight < 100%. ∗p ≤ 0.05 **e**, Visible signs of discomfort due to the infection were quantified daily and given a clinical score. **f**, Bar plots showing percentage of viral reads in the RNAseq datasets from mouse tissues of the infected mice. Only the lung tissue shows replication of the virus, with the levels going down by day4. **g**, Histological examination of lungs and assignment of scores based on severity of damage. **h**, Hematoxylin and eosin (H&E) staining of lung from non-infected mock controls, and SARS CoV-2 MA10-infected mice (wild-type and *Pcif1* KO). See Extended Data Fig. 5 for a more complete representation of this data. The days post infection (dpi) are indicated. The arrowheads mark lymphohistiocytic peribronchiolar inflammation, the stars mark intraluminal bronchiolar inflammation and necrosis, the asterisks mark interstitial lymphohistiocytic inflammation, and the arrows indicate vascular lesions such as vasculitis with surrounding lymphohistiocytic inflammation. Bars represent 100 µm (right column, higher magnification). **i**, Volcano plots showing gene expression changes in lungs between wildtype and *Pcif1* KO mice when infected with SARS CoV-2 MA. **j**, GO term analysis of gene expression alterations in the *Pcif1* KO lung.

To examine the relevance of PCIF1-mediated methylation for viral infection in an organismic model, we used the *Pcif1* KO mouse^9^. We used the mouse-adapted SARS CoV-2 (SARS CoV-2 MA10), which has a mutated spike protein that allows efficient attachment to the mouse ACE2 receptor^38^ (Fig. 4b). SARS CoV-2 MA10 causes acute lung injury in young BALB/c mice, while aged mice (1 year-old) display increased morbidity and almost complete lethality at high doses of 10^4^ and 10^5^ PFU^38^. In contrast, the infected C57BL6/J mice displayed very mild phenotypes^38^, and our *Pcif1* mutant is in this background^9^.

We intranasally infected control wildtype and *Pcif1* KO adult (8 weeks-old) mice of both sexes with a high dose (10^4^ or 10^5^ PFU) of SARS-CoV-2 MA10 during a 4-day period (Fig. 4c). Body weights were measured daily, and the animals were monitored for signs of obvious distress due to the viral infection, and clinical scores were assigned based on established criteria. To detect viral RNA by quantitative RT-PCR (qRT-PCR) or deep sequencing analyses, we collected throat swabs, and tissues (lung, nose, olfactory bulb and brain) at 2 days post infection (dpi) and 4dpi.

The infected control animals showed a decrease in body weight that started at 2dpi and continued until 4dpi. (Fig. 4d and Extended Data Fig. 4b). Although few of the *Pcif1* KO animals also displayed body weight reduction, this reduction was not consistent across individuals. The wildtype mice displayed symptoms like withdrawal, reduced cage activity, reduced feeding etc, already at day1, with 50% of the wildtype animals showing such symptoms and were given a clinical score of 0.5 (from a scale of 0-4) (Fig. 4e). By day2 this score climbed up to 1 for 50% of the animals, with the remaining animals being scored at 0.5. In contrast, the *Pcif1* KO mice seemed to handle the infection better with a delayed appearance of symptoms (scored at 0.5), which disappeared by day4 (Fig. 4e).

Detection of viral RNA by qRT-PCR shows higher levels in the lung at 2dpi, with no dramatic difference between the control and KO samples (Extended Data Fig. 4c-d). Deep sequencing analysis of lung tissue confirms this trend for higher viral RNA load at 2dpi compared to 4dpi (Fig. 4f) without any significant difference between the control and KO samples. There was also very high variability of detected viral loads between individual samples. Likewise, the histological analysis of the lungs^39^ of *Pcif1* KO and wildtype mice infected with SARS CoV-2 MA10 at 2 and 4 dpi, did not show major differences between the two groups with high variability among the samples (Fig. 4g). The most severely affected animals in both groups displayed a necrotizing bronchiolitis with peribronchiolar inflammation at day 2 post infection, and a mild to moderate interstitial pneumonia associated with occasional vascular lesions at day 4 post infection (Fig. 4h and Extended Data Fig. 5). However, there were big discrepancies regarding lesion severity within each group/sex/time point, and semi-quantitative assessment of lung pathology did not reveal significant differences between the obtained scores (Fig. 4g). As expected, RNAseq detected viral transcripts only in lungs and confirmed that it does not spread into other tissues, which correlates with observed mild phenotypes (Fig. 4f). This is in contrast to the severe phenotypes observed in the transgenic K18 model, where the hACE2 is driven from the cytokeratin-18 (K18) gene promoter, and ectopically expressed in multiple tissues^40^.

We also compared host gene expression in the different tissues of wildtype and *Pcif1* KO mice during the course of SARS CoV-2 infection (Fig. 4i-j and Extended Data Fig. 4e). In response to the infection in the lungs, we find several gene ontology categories enriched among the dysregulated genes only at 4dpi (Fig. j). At 2dpi we find elevated expression of some of genes associated with type I interferon response but with great variability between individuals and in both wild-type and mutant mice (Extended Data Fig. 4f-g) These results reveal that viral infection triggers interferon pathway activation in both control and *Pcif1* KO lung tissues, with no dramatic difference in viral RNA levels.

Newly produced full-length SARS CoV-2 viral genomic RNA and the sub-genomic RNAs are m^7^G-capped by the combined activity of several viral proteins^41^. The 5′ triphosphatase NSP13^42^, guanylyltransferase (GTPase) activity of the nidovirus RdRp-associated nucleotidyltransferase (NiRAN) domain^43,44^ of NSP12 (which is also the RNA-dependent RNA polymerase, RdRp), the *N^7^*-guanine methyltransferase of NSP14^45^ and the 2′-*O*-methyltransferase NSP16^34,37^. This results in the viral RNAs getting the cap1 structure, just like the mammalian host mRNAs. Despite this apparent self-sufficiency in terms of capping enzymes, here we show that the virus hijacks the host nuclear cap-dependent *N^6^*-adenosine methyltransferase PCIF1 to catalyse m^6^Am modification of the viral RNA caps. During viral infection, the long genomic RNA constitutes only a tiny fraction of the total RNA pool, with the sub-genomic RNAs dominating. Given that they both have a common 5′ leader sequence (76 nt)^10^, and the read-length in our m^6^A-IPseq dataset is in the range of 20-40 nt, we are unable to pin the reads to any particular RNA species. We propose that interaction of PCIF1 with either NSP12 (the RdRp) or other components of the viral capping machinery may recruit it to the nascent viral RNA for coordinated capping and m^6^Am installation. In addition, we report internal m^6^A modifications at multiple sites (Fig. 2a and 3a), as previously reported by others^28–31^. These are mediated by the host nuclear METTL3-METTL14 complex.

The loss of m^6^Am cap methylation in the *Pcif1* KO mice does not have any impact on viability or fertility^9^. However, levels of RNAs with a TSS adenosine are moderately reduced in some specific mutant tissues like the testis. In this study we find that SARS CoV-2 accumulates at a lower level (Fig. 3d) and supports reduced viral replication (Fig. 4a), in the *PCIF1* KO human cell cultures, but did not register a similar effect in the mouse *Pcif1* KO lung (Fig. 4f). The larger question is what selective advantage the m^6^Am modification provides to some coronaviruses like SARS CoV-2 and hCoV-229E. Our cell culture experiments point to a role for the 5′-terminal adenosine (Fig. 3f) and its m^6^Am methylation in supporting viral replication (Fig. 4a). In the case of vesicular stomatitis virus (VSV) infection, viral cap m^6^Am methylation by PCIF1 aids to attenuate host innate immune response^46^, but we did not observe such an effect in our SARS CoV-2-infected cell culture (Extended Fig. 3) and mouse (Extended Fig. 4) models. While our mouse experiments indicate a positive effect for the host in the absence of *Pcif1*(Fig. 4e), differences in viral replication or RNA levels (Fig. 4f) or innate immune responses were not apparent (Extended Data Fig. 4f-g). The big variability between individual samples (and of sampling tissue parts) might be a factor here. Future studies using the sensitive K18 mouse model^40^ might allow us to examine the true in vivo impact of loss of *Pcif1*. In summary, we show that m^6^Am modification on viral RNA cap is critical for maintaining viral RNA levels and for sustaining robust viral replication in cell cultures, and that absence of *Pcif1* in mice results in milder clinical outcomes after infection. This put PCIF1 and m^6^Am as new candidates to target in anti-viral therapies, especially given the lack of any serious phenotypes in the KO mice^9^.

## Methods

### Animal Work

The *Pcif1* KO mice were generated at the Transgenic Mouse Facility of University of Geneva for an earlier study^9^. The mice were bred in the Animal Facility of Sciences III, University of Geneva. The use of animals in research at the University of Geneva is regulated by the Animal Welfare Federal Law (LPA 2005), the Animal Welfare Ordinance (OPAn 2008) and the Animal Experimentation Ordinance (OEXA 2010). The Swiss legislation respects the Directive 2010/63/EU of the European Union. Any project involving animals has to be approved by the Direction Générale de la Santé and the ethics committee of the Canton of Geneva, performing a harm-benefit analysis of the project. Animals are treated with respect based on the 3Rs principle in the animal care facility of the University of Geneva. We use the lowest number of animals needed to conduct our specific research project. Discomfort, distress, pain and injury is limited to what is indispensable and anesthesia and analgesia is provided when necessary. Daily care and maintenance are ensured by fully trained and certified staff. All animals were housed 3-5 per cage and maintained on a 12-hour light/dark cycle, with water and food available ad libitum. The use of wildtype control and *Pcif1* KO mice for SARS CoV-2 MA infection studies in the BSL3 facility was approved by the Canton of Bern (xxxxxx).

### *Pcif1* mutant mouse line

The *Pcif1* knockout mice are previously described^9^. Genotyping protocols are also described in detail in that study.

### Cell lines

The control wildtype Human Embryonic Kidney 293T (HEK293T) cells, and those lacking *PCIF1*^8^, and were a kind gift of Eric Lieberman Greer (Harvard Medical School). HEK293T cells are not efficiently infected by SARS-CoV-2, therefore we introduced an expression cassette for the cell-surface receptor for the virus, the human Angiotensin-converting enzyme 2 (ACE2). Lentiviruses carrying the expression cassette were prepared and used to infect the HEK293T cell lines. The resulting HEK293T (ACE2) cells were confirmed to stably express *ACE2*. African green monkey kidney-derived Vero E6 cells and human liver cell line Huh7 were obtained from commercial sources.

### SARS CoV-2 viruses used in this study

A clinical isolate of SARS CoV-2 from Munich was used for the 5′ RACE experiment to map the 5′ end of the viral RNA. All other experiments used a recombinant preparation^32^ of the human SARS CoV-2 that is identical in sequence to the index SARS-CoV-2 (Wuhan-Hu-1). To generate recombinant coronaviruses a T7-transcribed 30 kb viral RNA preparation is used for electroporation into mammalian cells (BHK-21 cells expressing the SARS CoV nucleocapsid protein; BHK-S-N). Following electroporation, the BHK-S-N cells are co-cultured with SARS-CoV-2-susceptible cells, such as Vero E6 cells, to allow for virus spread and propagation in the initial culture and to obtain the recombinant SARS-CoV-2 in the cell culture supernatant^47^. Mice were infected with the SARS CoV-2 MA virus^38^ which has a mutated spike allowing attachment to the mouse ACE2 receptor.

### Antibodies

The polyclonal rabbit anti-m^6^A (Synaptic Systems; cat. no. 202003) for m^6^A-IP-seq.

### Histology of SARS CoV-2 MA10-infected mouse tissues

The left lung was collected during necropsy at 2 and 4 days post infection (dpi) with the mouse-adapted SARS CoV-2 MA10 virus^38^ and immersed in 10% neutral-buffered formalin. Following fixation, tissues were embedded in paraffin, cut at 4 μm, and stained with haematoxylin and eosin (H&E) for histological evaluation. Lung tissue pathology was scored according to a previously published scoring scheme^39^.

### Poly(A) RNA and SARS CoV-2 RNA purification

#### Poly(A)+ RNA purification

SARS CoV-2 infected cells total RNAs was isolated using TRIZOL reagent. The poly(A) RNA was purified using Dynabeads mRNA Purification Kit (Invitrogen, cat. no. 61006). All the steps were performed according to the protocol provided by the manufacturer. Briefly, 225 µg of RNA was diluted in RNase-free water to a final volume of 300 µL. RNA was then heated to 65°C for 3 minutes and placed on ice. In the meantime, 600 µL of resuspended Dynabeads Oligo (dT)_25_ beads for each sample were transferred to fresh RNase-free tubes. The tubes were placed on a magnetic rack for 2 minutes, and the supernatant was removed. The beads were resuspended in 300 µL of Binding Buffer, placed on a magnetic stand, and the supernatant was removed. Next, the beads were resuspended in 300 µL of Binding Buffer and then were mixed with the RNA (1:1 volume ratio). The RNA was incubated with the beads for 5 minutes at room temperature with rocking. Then, tubes were placed on the magnetic stand and supernatant was removed. Beads were washed twice with 600 µL Washing Buffer B. Finally, the beads were resuspended in 40 µL of water, heated to 75°C for 2 minutes and immediately placed on the magnet. The supernatant with eluted mRNA was transferred into fresh RNase-free tubes. The eluted poly(A)+ RNA was either used immediately or stored at -80°C, to avoid RNA degradation.

#### SARS Cov-2 RNA purification

Total RNA (600 µg) from SARS CoV-2 infected Vero E6 cells was used to purify poly(A)+ RNAs using Poly(A)Purist MAG kit (Thermo Fischer Scientific, cat. No. AM1922) according to manufacturer’s instructions. The poly(A)+ RNA was cleaned with RNA Clean and Concentrator-5 (Zymo research, R1016) following the kit’s total RNA cleanup protocol and eluted in 15 µL water. SARS CoV-2 RNA was purified from poly(A)+ RNA using biotinylated antisense DNA oligo against the SARS CoV-2 leader sequence (Table S1). Briefly, 5.0 µg of poly(A)+ RNA was mixed with 5µL hybridization buffer (10 mM Tris-HCl (pH 7.5), 1 mM EDTA, 2 M NaCl), 1 µL of antisense oligo (RRoligo1227, RRoligo1228, RRoligo 1230 and RRoligo1231, 100µM each) and RiboLock RNase Inhibitor (Thermo Scientific, EO0381) in a 20 µL reaction volume. The reaction content was thoroughly mixed with a 20 µL pipette, briefly centrifuged and incubated at 68°C for 10 minutes, then cooling to 37°C in a thermomixer to hybridize the antisense oligos with RNAs (Table S1). In the meantime, 100 µL MyOne C1 magnetic beads suspension for each sample (Thermo scientific, Cat No. 65002) was taken into a fresh RNase-free tube and washed twice with 100 µL bead resuspension buffer (0.1 M NaOH, 0.05 M NaCl), once with 100 µL wash buffer (10 mM Tris-HCl (pH 7.5), 0.15 M LiCl, 1 mM EDTA). During each wash, beads were mixed with the buffer by a gentle vortex, placed on the magnetic rack for a minute, and removed supernatant. Finally, beads were resuspended in 160 µL of depletion buffer (10 mM Tris-HCl (pH 7.5), 1 mM EDTA, 1 M NaCl). The hybridized RNA-antisense oligo mix was transferred to MyOne C1 beads containing tubes, mixed well with the pipette and incubated at 37°C for 15 minutes. The tubes were transferred to a preheated heat block at 50°C and incubated for another five minutes. At the end of the incubation, tubes were quickly transferred to the magnetic rack, and while tubes still are on the magnetic rack, the supernatant was carefully removed without touching the beads. The beads were washed three times with 100 µL wash buffer. After three washes, the beads were resuspended in 20 µL elution buffer (10mM Tris-HCl pH 7.0), incubated at 75°C in a preheated heat block for 2 minutes, quickly transferred to the magnetic rack, and the supernatant was collected. The purified RNA was used for RNA-seq to determine SARS CoV-2 RNA enrichment and RNA mass spectrometry to identify modified bases.

### Quantification of RNA modifications using LC-MS/MS

Total RNA was isolated by Trizol extraction from mock-infected or SARS CoV-2-infected Vero E6 or HEK293T ACE2 wildtype or *PCIF1 KO* cells. Poly(A)+ RNA was purified from total RNA using Dynabeads mRNA Purification Kit (Invitrogen, cat. no. 61006). The poly(A)+ RNAs from Vero cells were further purified with SARS CoV-2 specific biotinylated antisense DNA oligos to enrich viral RNAs. The purified RNAs were divided into two parts: one part was hydrolyzed to ribonucleosides by 20 U Benzonase® Nuclease (Santa Cruz Biotech, cat. no. sc-202391) and 0.2 U Nuclease P1 (Sigma, cat. no. N8630-1VL) giving information on modifications on the entire RNA (cap+internal), whereas the other part was treated with nuclease P1 only and gives modification information only on the body of the RNA (internal only), in 10 mM ammonium acetate pH 6.0 and 1 mM magnesium chloride at 40°C for one hour. After that, ammonium bicarbonate to 50 mM, 0.002 U phosphodiesterase I and 0.1 U alkaline phosphatase (Sigma) were added and incubated further at 37°C for 1 h. The hydrolysates were mixed with 3 volumes of acetonitrile and centrifuged (16,000 x g, 30 min, 4°C). The supernatants were dried and dissolved in 50 µL water for LC-MS/MS analysis of modified and unmodified ribonucleosides. Chromatographic separation was performed using an Agilent 1290 Infinity II UHPLC system with an ZORBAX RRHD Eclipse Plus C18 150 x 2.1 mm ID (1.8 µm) column protected with an ZORBAX RRHD Eclipse Plus C18 5 x 2.1 mm ID (1.8 µm) guard column (Agilent). The mobile phase consisted of water and methanol (both added 0.1% formic acid) run at 0.23 mL/min. For modifications, starting with 5% methanol for 0.5 min followed by a 2.5 min gradient of 5-15% methanol, a 3 min gradient of 15-95% methanol and 4 min re-equilibration with 5% methanol. A portion of each sample was diluted for the analysis of unmodified ribonucleosides, which was chromatographed isocratically with 20% methanol. Mass spectrometric detection was performed using an Agilent 6495 Triple Quadrupole system, monitoring the mass transitions 268.1-136.1 (A), 284.1-152.1 (G), 244.1-112.1 (C), 245.1-113.1 (U), 296.1-150.1 (m^6^Am), 282.1-150.1 (m^6^A and m^1^A), 285.1-153.1 (*d*_3_-m6A), 282.1-136.1 (Am), 296.1-164.1 (m^6,6^A), 283.1-151.1 (m^1^I), 298.1-166.1 (m^7^G), 312.1-180.1 (m^2,7^G), 326.1-194.1 (m^2,2,7^G), 258.1-126.1 (m^3^C and m^5^C) in positive electrospray ionization mode, and 267.1-135.1 (inosine, I) and 243.1-153.1 (pseudouridine, Ψ) in negative electrospray ionization mode. Modifications detected in a mock control (containing only the hydrolytic enzymes) were subtracted from modifications detected in the RNA samples. In general, the mock control contained at least 1000-fold less RNA than the RNA samples and gave negligible background in the modification analyses.

### 5’ RACE to identify the TSS nucleotide of SARS CoV-2 RNAs

Total RNA from Vero E6 cells infected with SARS Cov-2 was purified and 2.0 µg of the RNA was decapped with five units of Cap-Clip Acid Pyrophaosphatase (Cat. No. C-CC15011H, Cellscript) in a 20 µL reaction at 37°C for 30 minutes. The RNA was cleaned up with RNA Clean and Concentrator-5 (Zymo research, R1016) and eluted in 9.5 µL water. The decapped RNA was ligated with an RNA adapter in a total 25 µL reaction (RNA 9.25 µL, 10x T4 RNA ligase buffer 2.50 µL, 5 µM 5′ Adapter in 3.0 µL (15 pmol total), RNasin plus 1.0 µL (40units), T4 RNA ligase 2.0 µL (NEB, M0204S), 50% PEG8000 6.25 µL, 25 mM ATP 1.0 µL) overnight at 16°C. The next day, RNA was purified with RNA Clean and Concentrator-5 (Zymo research, R1016) following the long RNA purification protocol and subjected to first-strand cDNA synthesis using 0.5 µg (1 µL, 50 µM stock) primer pWhSF-ORF1a-R18-655 (Supplementary Table 1) binding to the ORF1a, and superscript RTII (ThermoFisher Scientific, Cat. 18064022). The reverse-transcription (RT) reaction was performed at 25°C for 5 minutes and followed by 42°C for one hour. The RT reaction was inactivated at 65°C for 20 minutes. After inactivation of the RT reaction, 1 µL RNase H was added to the reaction mix and incubated further for 20 minutes at 37°C. The cDNA was purified using High Pure PCR Product Purification Kit (Roche Cat. No. 11 732 668 001) as indicated below. Add 125 ul Binding Buffer (green cap) to 25 ul of the first-strand cDNA reaction and mix well. Combine the High Pure Filter Tube and the Collection Tube and pipet sample into the upper reservoir. Centrifuge for 30 sec at 6,000 to 8,000 × g in an Eppendorf centrifuge. Discard the flow through liquid from collection tube and add 500 ul Wash Buffer to the upper reservoir of the Filter Tube. Centrifuge for 30 sec at 6,000 to 8,000 × g in an Eppendorf centrifuge. Make sure that the Filter Tube has no contact with the surface of the Wash Buffer flow through. Reinsert the Filter Tube into the same Collection Tube. Discard the flow through and add 200 ul Wash Buffer to the upper reservoir of the Filter Tube. Centrifuge at least 2 min at maximum speed (approximately 13,000 × g) in an Eppendorf centrifuge. This additional washing step with reduced buffer volume ensure optimal purity and the complete removal of residual wash buffer from the glass fiber fleece. Carefully remove the filter tube from collection tube and insert it into a sterile 1.5 ml microcentrifuge tube. Add 50 ul Elution Buffer (Vial 3) to the Filter. Centrifuge for 30 sec at 6,000 to 8,000 × g in an Eppendorf centrifuge and save the elute, which contains cDNA. Primer sequences are given in Supplementary Table 1.

The purified cDNA was used for ***1^st^ PCR amplification*** in a 20 µL final reaction volume as follows: 10 µL HotStarTaq Master mix 2x, 1 µL pWhSF-ORF1a-R18_655 of 10 μM, 1 µL of RNA_PCR_Primer (RP1) of 10 μM, 2 µL cDNA and 6 µL water. PCR cycle conditions: Polymerase activation with first cycle at 95°C, 15 min; cycling touch-down 15 cycles of 94°C for 30 sec, 65-50°C (annealing temperature is gradually brought down to 50°C) for 1 min, 72°C for 1 min; cycling with 25 cycles of 94°C for 30 sec, 50°C for 1 min, 72°C for 1 min; final extension with one cycle of 72°C for 10 min; storage at 4°C.

The PCR product from 1^st^ PCR amplification was used to ***reamplify the DNA***. A total of 50 µL reaction volume: 25 µL HotStarTaq Master mix 2x, 3 µL pWhSF-5ut-R17_273 of 10 μM, 3 µL of RNA_PCR_Primer (RP1) of 10 μM, 2 µL 1^st^ PCR product and 17 µL water. The PCR cycle was same as the 1^st^ PCR. The PCR product was analysed on 2% agarose gel and the correct size band was purified and sequenced by Sanger sequencing with the primer pWhSF-5utr-R17-273. Primer sequences are given in Supplementary Table 1.

### Preparation of RNA libraries

#### RNA sequencing of purified SARS CoV-2 RNA

After a two-step purification (polyA+ and SARS CoV-2 specific antisense oligos) RNA were subjected to deep sequencing for quantifying viral RNA in our purified samples, and for RNA mass spectrometry (Fig. 1). Briefly, 40 ng of RNA was used to prepare the libraries using True-Seq kit from Illumina according to manufactures protocol. The ribodepletion step was omitted and the RNAs were directly subjected to fragmentation and steps thereafter. The libraries were sequenced from single end for 150 cycles on Illumina HiSeq4000 instrument.

#### m^6^A-IPseq to map m^6^A on SARS CoV-2 RNA

To map the location m^6^A methylation on SARS CoV-2 RNA genome, we carried out m^6^A-IPseq using polyA+ RNA from different cell lines infected with SARS CoV-2. The poly(A)+ RNAs (∼3.0 µg) was fragmented with 2 µL of fragmentation reagent (AM8740, Thermo Fisher Scientific) in a final volume of 20 µL in a PCR tube. The reaction mix was incubated at 75°C for 12 minutes in a PCR machine. The tube was then transferred on ice immediately, and the reaction was stopped by adding 2.2 µL of stop solution provided with fragmentation reagent. Denaturing urea-PAGE confirmed that most of the RNA fragments were in size range of 20-80 nts. One-tenth (10%) of fragmented RNA from each sample was kept aside as input, while the remainder was subjected to immunoprecipitation (IP).

The m^6^A immunoprecipitation was performed as described (Ke et al., 2015). Briefly, Protein A Dynabeads were washed once in PXL buffer (1× PBS, 0.1% SDS, 0.5% sodium deoxycholate, 0.5% NP-40) followed by pre-treatment with BSA (final concentration 1 µg/ µL) in 200 µL PXL buffer for 45 minutes at RT. BSA pre-treated beads were then conjugated with m^6^A rabbit polyclonal antibody (20 µg; Synaptic Systems, catalog no. 202003) in 200 µL PXL buffer supplemented with 4 µL of Ribolock RNAse inhibitor (EO0381, ThermoFisherScientific) for one hour at RT on a rotating wheel. Dynabeads were washed twice with PXL buffer and then resuspended in 400 µL of PXL buffer and 5 µl of Ribolock RNAse inhibitor. Fragmented RNA was added to the beads and incubated at 4°C for 2 hours on a rotating wheel. After two hours incubation, the beads were washed: twice by ice-cold Nelson low-salt buffer (15 mM Tris at pH 7.5, 5 mM EDTA), once by ice-cold Nelson high-salt buffer (15 mM Tris at pH 7.5, 5 mM EDTA, 2.5 mM EGTA, 1% Triton X-100, 1% sodium deoxycholate, 0.1% SDS, 1 M NaCl), once by ice-cold Nelson stringent wash buffer (15 mM Tris at pH 7.5, 5 mM EDTA, 2.5 mM EGTA, 1% Triton X-100, 1% sodium deoxycholate, 0.1% SDS, 120 mM NaCl, 25 mM KCl), and last by ice-cold NT-2 buffer (50 mM Tris at pH 7.4, 150 mM NaCl, 1 mM MgCl2, 0.05% NP-40). Antibody bound RNAs were eluted twice by incubating the beads with 200 µl of 0.5 mg/mL *N^6^*-methyl adenosine (Sigma-Aldrich; M2780) in NT2 buffer for one hour at 4°C. The eluted RNAs were precipitated overnight at - 20°C with ethanol and glycogen and dissolved in 35 µL RNase-free water. The input and IP RNAs were first 3ʹ end dephosphorylated with T4 PNK (NEB; M0201S, 10 U/mL) in the absence of ATP at 37°C for 45 minutes (40 µL reaction: 35.5 µL RNA, 4 µL 10X T4 PNK buffer, 0.5 µL of T4 PNK) followed by phosphorylation of 5ʹ end (50 µL reaction: 40 µL dephosphorylated RNA, 6.5 µL water, 1 mL RNasin, 0.5 µL 100 mM ATP, 1 µL 10X T4 PNK buffer 1 µL T4 PNK) at 37°C for 45 minutes. RNAs were phenol chloroform-extracted, ethanol precipitated and resuspended in 6 mL of RNase free water.

The input RNA fragments and the immunopurified RNAs, after the phosphorylation step, were directly used for strand-specific library preparation (barcoded at 3′ end) using NEBNext® Multiplex Small RNA Library Prep Set for Illumina® (NEB; catalogue No. E7560L) following manufacturer’s instructions. The libraries were resolved on 3% high-resolution MethaPhor agarose (Lonza; catalog. No. 50180) gels in 1X TAE buffer at 70 V. Fragments in the size-range of ∼150-250 bp were gel-extracted with the use of MinElute Gel Extraction Kit (Qiagen; cat No. 28604). Multiple libraries with different barcodes (at the 3′ end) were mixed in equimolar ratios and sequenced with the NextSeq Illumina Platform (EMBL Gene Core facility, Heidelberg). The maximum sequencing length was 75 nucleotides. The list of sequencing libraries generated is provided in Supplementary Table 2.

### Computational analyses

#### Quantification of SARS CoV-2 RNA from RNA sequencing

Sequenced reads from Poly(A)+ isolated RNAs of Vero cells infected with SARS CoV-2 RNA (samples RR947:RR949) and those after SARS-CoV-2 two step affinity purification (samples RR938:RR940) were aligned to the green monkey (ChlSab1.1;Ensembl release 95) and SARS CoV-2 (MT108784) genome using STAR 2.7.0a (parameters: --outFilterType BySJout --limitOutSJcollapsed 50000000 --limitIObufferSize 1500000000). Proportion of the mapped and non-mapped reads was plotted (Fig. 1b).

#### Plotting the quantification of RNA modifications using LC-MS/MS

The heatmap was plotted in R 3.6.3 (https://www.r-project.org/) using gplots::heatmap.2 (Fig. 1c) and barplots were made for selected modifications (Fig. 1e, Extended Data Fig. 1a).

#### Analysis of m^6^A-IP seq libraries

The reads were sorted into individual libraries based on the barcodes and clipped using cutadapt 2.0 (parameters: -a AGATCGGAAGAGCACACGTCT -m 15 -e 0.2 -O 4 -q 10 --match-read-wildcards; parameter -A GATCGTCGGACTGTAGAACTCTGAAC used in addition for pair-end sequencing). The clipped reads were aligned using STAR 2.7.0a (parameters: --outFilterType BySJout --limitOutSJcollapsed 50000000 --limitIObufferSize 1500000000). The samples from SARS-CoV-2 infected Vero cells were aligned to SARS CoV-2 (MT108784) and green monkey (ChlSab1.1;Ensembl release 95) genome. The samples from hCoV-229E infected Huh7 cells were aligned to hCoV-229E (NC_002645.1) and human (GRCh38;Ensembl release 95) genome.

The samples from SARS CoV-2 infected HEK293T (ACE2) cells were aligned to SARS CoV-2 (MT108784) and human (GRCh38;Ensembl release 95) genome. The whole read coverages along the viral genomes were created using IRanges::coverage function in R. Similarly, the coverages were calculated only for the 5′ ends of the reads – only read1 was used in case of pair-end sequencing. The coverages were normalized to library sizes (rpm) and the read coverage IP enrichments were calculated by dividing the m^6^A-IP coverage by input coverage. Means of three replicas were plotted along whole genome or only the 5′ end part of the genome using whole read coverages or coverages of 5′ ends of the reads (Fig. 2a,b, 3a,b; Extended Data Fig. 2a,b, 3a). To compare the amount viral of m^6^Am, the 5’ read coverage enrichment was plotted for the 5′ adenosine (position 1) of the viral genome (Fig. 2c,3b, Extended Data Fig. 2,3a). Proportion of reads mapped to the viral genome was plotted from input samples to compare the viral RNA levels (Fig. 2d,3d, Extended Data Fig. 2d,3c).

#### Analysis of gene expression

Input samples from m^6^A-IP seq experiments or RNAseq samples were used to analyses the gene expression. The reads were mapped using SALMON 1.3.0 (parameters: -l A -p 10–gcBias–validateMappings) to the host transcriptome (GRCh38;Ensembl release 95 for human, GRCm38;Ensembl release 95 for mouse) and differential gene expression was analysed using DESeq2 1.25.10. Obtained adjusted p-values and log2 fold changes for individual genes were used to construct volcano plots using EnhancedVolcano function from EnhancedVolcano 1.3.5 package (Fig. 4i,Extended Data Fig. 3d,e,4e). The dysregulated gene lists were searched for enriched Gene Ontology biological process terms using ENRICHR (https://maayanlab.cloud/Enrichr/) and top ten enriched categories were shown (Fig. 4j, Extended Data Fig. 3d,e). For genes belonging to “Cellular response to type I interferon genes” (GO:0071357) and interferon genes, the heatmaps were constructed plotting the z-score of expression in individual samples using pheatmap_1.0.12 together with boxplots of the log2 fold changes (Extended Data Fig. 3f, 4f,g).

## Reporting Summary

Further information on research design is available in the Nature Research Reporting Summary linked to this article.

## Data availability

The data that support the findings of this study are available from the corresponding authors on request. The deep sequencing data is deposited with the Gene Expression Omnibus (GEO: GSEXXX). Raw data used in preparation of the figures will be deposited with Mendeley Data http://dx.doi.org/10.17632/hbn9vgbww5.1

## Acknowledgements

We thank Eric Lieberman Greer (Harvard Medical School) for PCIF1 KO HEK29T cells. We thank Pacal Gos for help with mouse work and Nicolas Roggli for scientific illustration. We thank the following facilities: iGE3 Genomics Platform, University of Geneva; the EMBL Genomics core facility. This work was supported by grants to R.S.P and V.T from the Swiss National Science Foundation: Project grant 310030B_201278 (to VT), Special Call on Coronaviruses 2020 (31CA30_196387) and NRP 78 Covid-19 2020 (4078P0_198473), and funding from the NCCR RNA & Disease (51NF40_182880). Work in the Pillai lab is supported by the Republic and Canton of Geneva.

## Author contributions

R.R.P lentiviral transduction of cell lines, m^6^A-IPseq and library preparations and RACE; N.E all virus infection experiments in the BSL3 lab; D.H computational analysis; T.B, B.S.T, H.S designed recombinant viruses and carried out RACE; E.D Western analysis; C.B.V conducted RNA mass spectrometry; I.B.V histological analyses; R.S.P, V.T and S.A.L supervised the study and wrote the manuscript with input from everyone.

## Additional information

Extended data is available for this paper at

Supplementary information is available for this paper at

Correspondence and requests for materials should be addressed to R.S.P and V.T.

---

**Extended Data Fig. 1.**
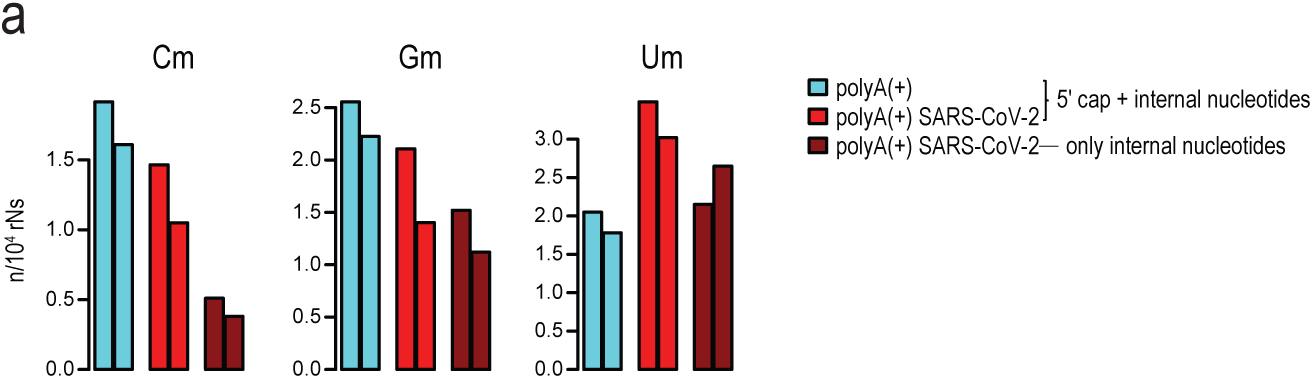
Ribose RNA methylations of SARS CoV-2. **a**. PolyA+ RNA or affinity-purified SARS CoV-2 RNA was subjected to mass spectrometry. Bar plots show abundance (number of modified nucleotides/10^4^ nucleotides) of some of the modifications. Duplicate biological replicates were tested.

**Extended Data Fig. 2.**
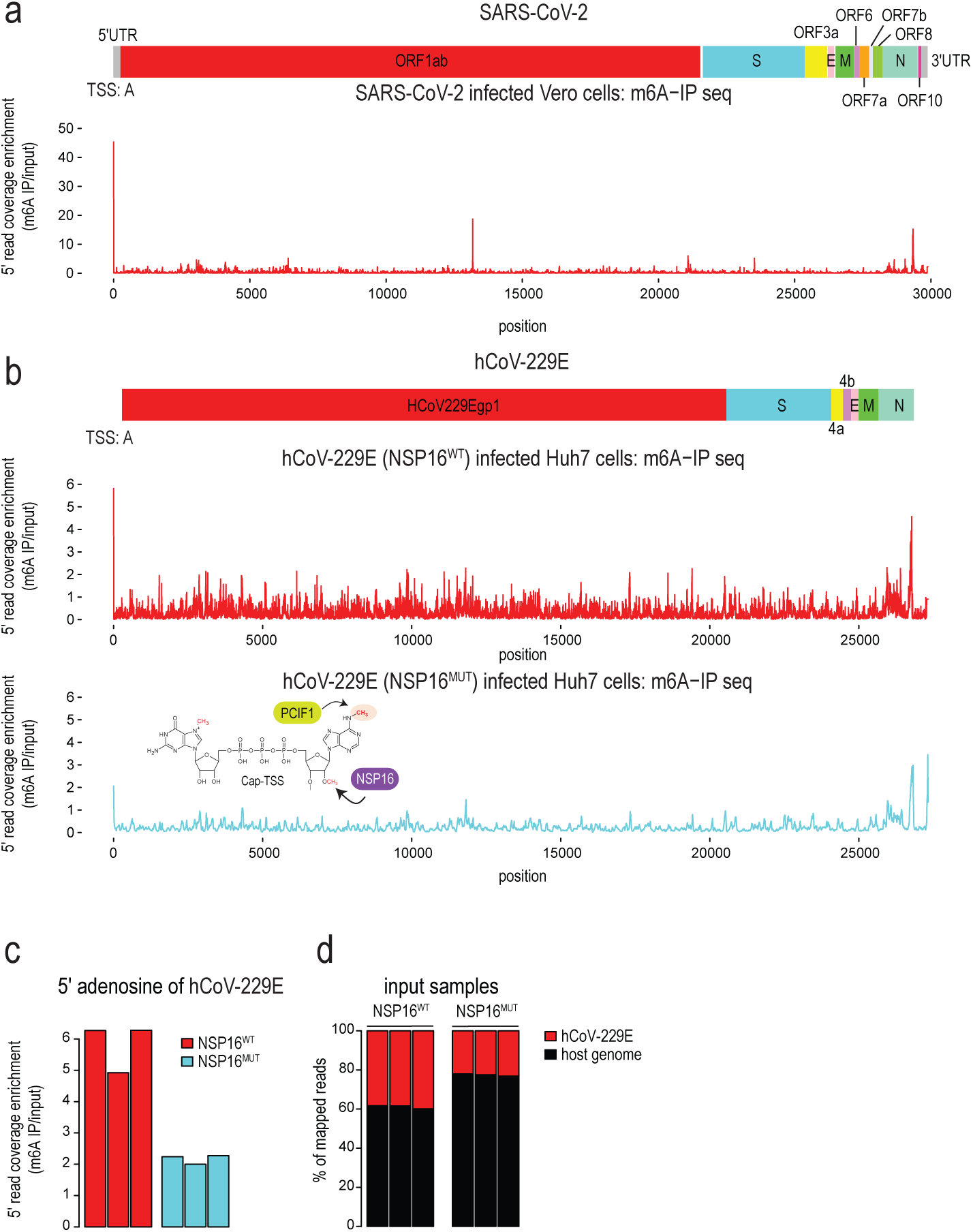
Virus-encoded NSP16 is required for efficient m^6^Am methylation of coronavirus cap structure. **a**, Mapping the reads from infected Vero E6 cells to SARS CoV-2 genome and plotting the enrichment of their 5′ end coverage in m^6^A-IPseq libraries (three replicas). The prominent 5′ end peak corresponds to m^6^Am as the coronavirus has a cap1 structure (Am). **b**, Mapping of m^6^A on the common cold-causing human coronavirus hCoV-229E using m^6^A-IP seq (three replicas) with RNA from infected Huh7 cells. The prominent 5′ end peak corresponding to m^6^Am is reduced when cells are infected with a virus mutant for the virus-encoded *NSP16* that catalyzes 2′-*O*-methylation of the TSS nucleotide (Am) (three replicas). **c**, Bar plot showing enrichment in m6A-IPseq libraries for the reads starting at the 5′ adenosine, which demonstrates a reduction in m6Am in the NSP16 mutant hCoV-229E. Three replicas are shown separately. **d**, Bar plot showing proportion of reads mapping to the viral and host genome which demonstrates the reduced viral RNA levels when cells are infected with the NSP16 mutant. Three replicas are shown separately.

**Extended Data Fig. 3.**
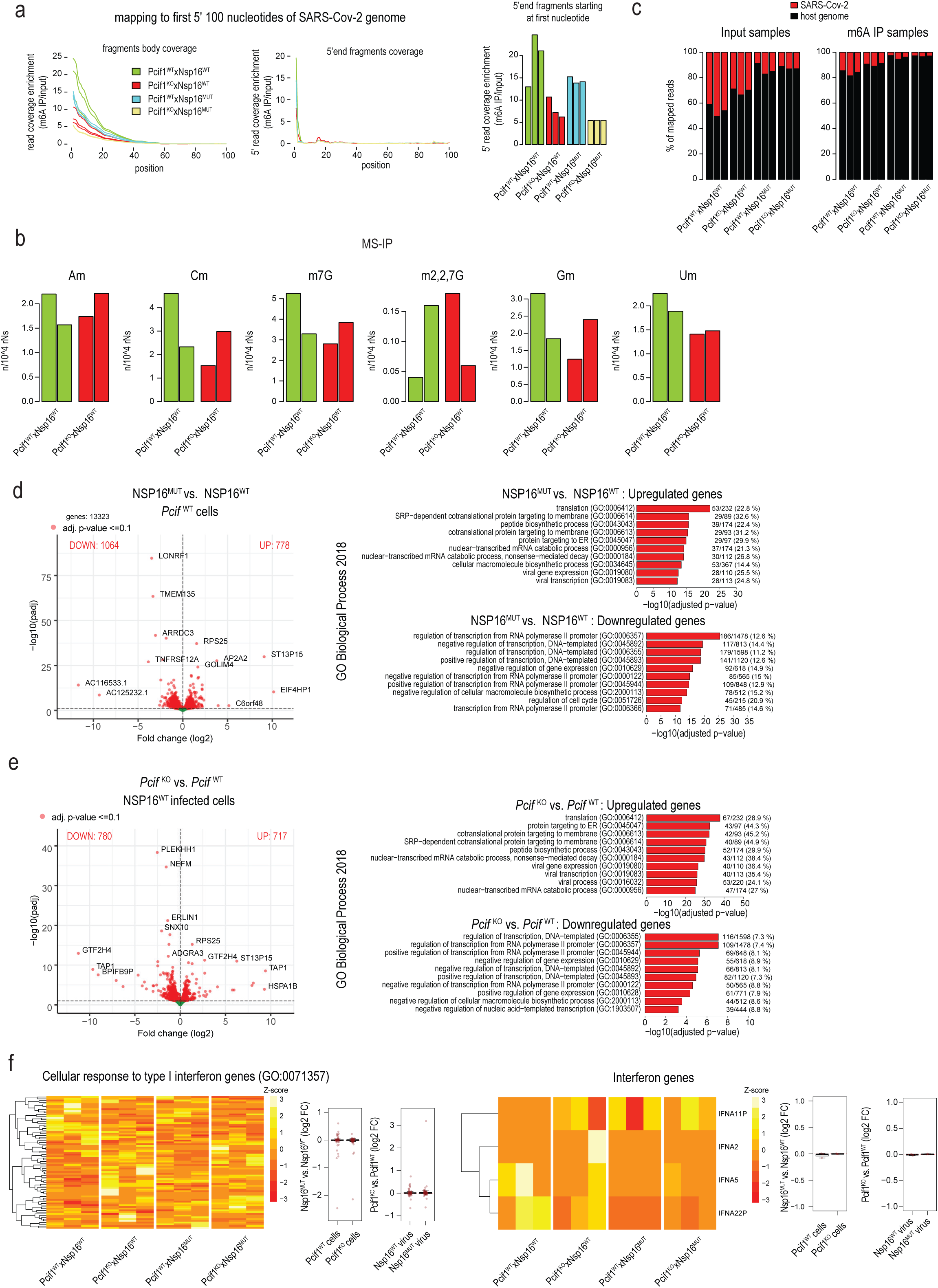
m^6^Am cap methylation on SARS CoV-2 is catalyzed by host PCIF1. **a**, Wildtype or *PCIF1* KO human HEK293T ACE2 cells were infected with SARS CoV-2 having wild-type or mutated NSP16. Enrichment in m^6^A-IPseq libraries is plotted for coverages of whole reads or of their 5′ ends. Only 5′ region of the viral genome is shown. Bar plot shows enrichment in m6A-IPseq libraries for the reads starting at the 5′ adenosine. Three replicas are shown separately. **b**, RNA mass spectrometry of polyA+ RNA from wildtype or *PCIF1* KO HEK293T ACE2 cells. Bar plots show some of the RNA modifications detected. While Fig. 3 shows that there is the expected complete absence of m^6^Am in the KO cells, these bar plots show that other modifications are not dramatically affected. Experiments done in duplicates. **c**, Bar plot showing abundance of viral RNA as a percentage of viral mapped reads among the total reads mapped to the viral or host genome. d, Volcano plots showing gene expression changes between *Pcif1* WT cells infected with SARS CoV-2 having wilt-type or mutated NSP16. Top ten GO terms (Biological process) enriched among upregulated and downregulated genes are shown on the right. **e**, Volcano plots showing gene expression changes between *Pcif1* KO and WT cells infected with SARS CoV-2 having wilt-type NSP16. Top ten GO terms (Biological process) enriched among upregulated and downregulated genes are shown on the right. **f**, Heatmaps show the expression of type I interferon pathway genes and interferon genes in individual samples. Boxplots show expression changes between *Pcif1* KO and WT cells or cells infected by SARS CoV-2 having wilt-type or mutated NSP16. Dots represent the individual genes.

**Extended Data Fig. 4.**
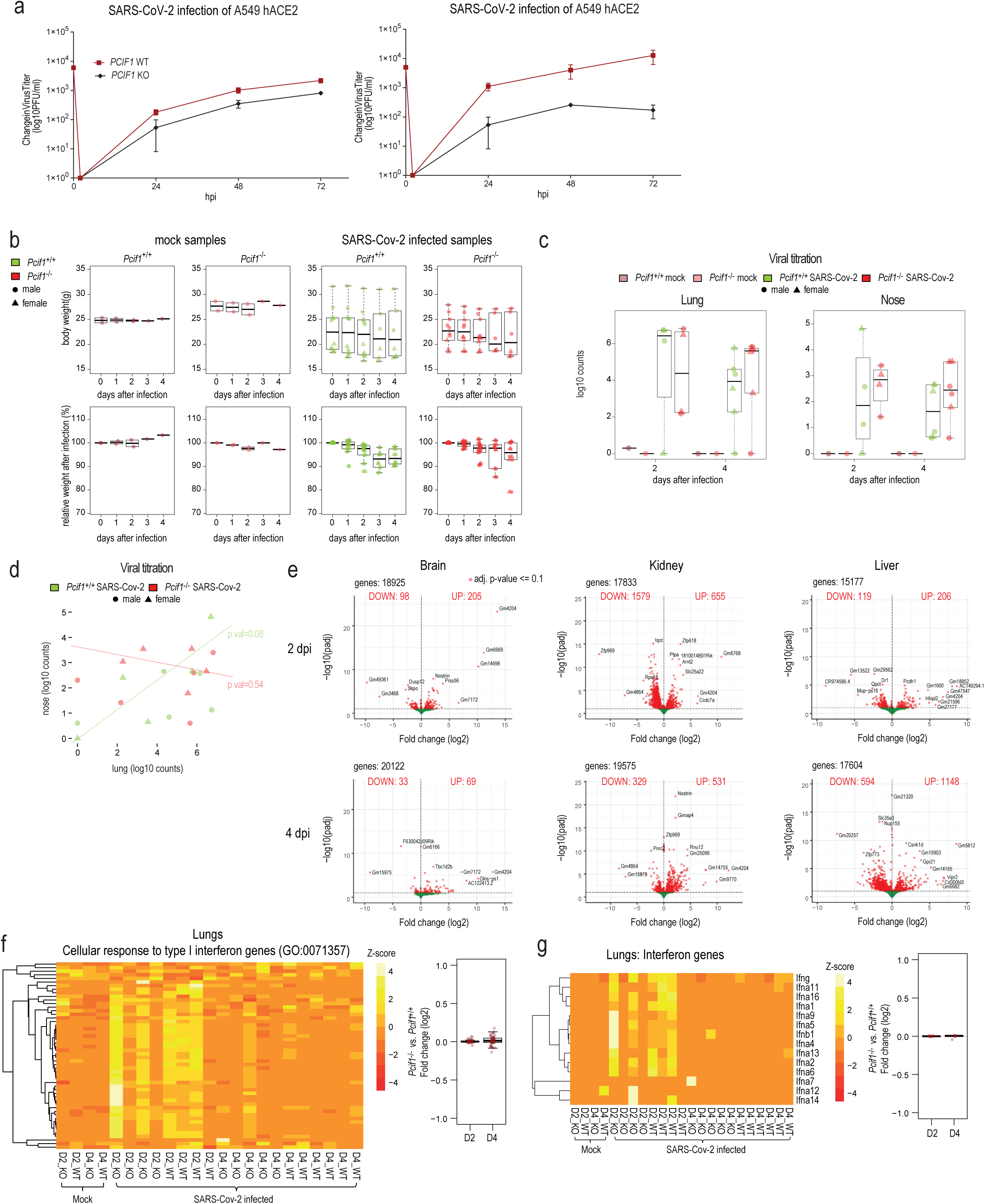
Reduced SARS CoV-2 replication in PCIF1-depeleted cells and gene expression changes in mice infected with SARS CoV-2. **a**, Infection of A549 ACE2 WT and PCIF1-depleted (Mut) cells with SARS CoV-2. Viral titres estimated by plaque assays. This is in addition to the experiment presented in Fig. 4a. **b**, A mouse-adapted version of human SARS CoV-2 coronavirus with mutations (SARS CoV-2 MA) in the spike that allows attachment to the mouse ACE2 receptor was used for mouse infection studies. Either wildtype (n=10) or *Pcif1* KO (n=10) mice were infected with SARS CoV-2 MA at day 0 and followed for a period of 4 days. Mock-treated animals were left uninfected. The measured body weight and relative change in weight after infection are shown. **c**, Viral load in the lung and nose buds were quantified by RT-PCR at day2 and day4. **d**, Comparison of viral loads in the nose buds and in the lungs does not show significant correlation. **e**, Volcano plots showing gene expression changes between SARS CoV-2 MA infected wildtype and *Pcif1* KO mice. Gene expression was analyzed in multiple tissues at day2 and day4. None of these tissues shows presence of the viral RNA, which was detected only in lungs. **f**, Heatmap shows the expression of type I interferon pathway genes in the lungs of uninfected (mock) mice and mice infected with SARS CoV-2 MA. Expression in individual samples is plotted. Note the increased expression of the type I interferon pathway genes at day2 post infection (D2) in both the infected wildtype and *Pcif1* KO mice, and this subsides by day4 (D4). Box plot shows that there is no change in the type I interferon pathway gene expression due to loss of *Pcif1*. **g**, Heat-map focusing only on interferon gene expression in the lung of mock uninfected and infected mice. Box plot shows no impact due to loss of *Pcif1*.

**Extended Data Fig. 5.**
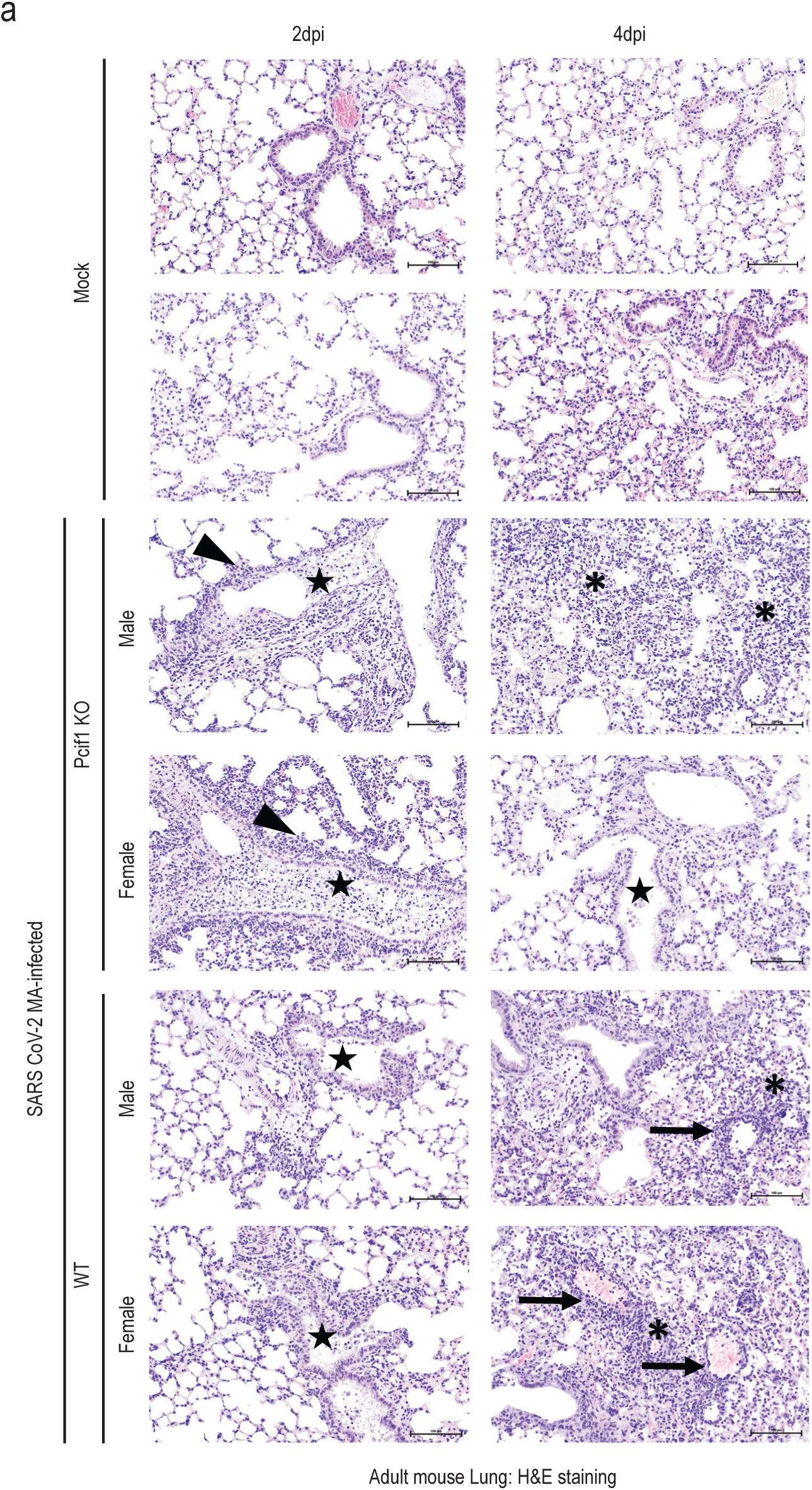
Histopathological analysis of *Pcif1* KO and wildtype mouse lung tissue infected with SARS CoV-2 MA10. **a**, Hematoxylin and eosin (H&E) staining of lung from non-infected mock controls, and SARS CoV-2 MA10-infected mice (wild-type and *Pcif1* KO). The days post infection (dpi) are indicated. The arrowheads mark lymphohistiocytic peribronchiolar inflammation, the stars mark intraluminal bronchiolar inflammation and necrosis, the asterisks mark interstitial lymphohistiocytic inflammation, and the arrows indicate vascular lesions such as vasculitis with surrounding lymphohistiocytic inflammation. Bars represent 100 µm (right column, higher magnification). A part of this data is also presented in Fig. 4h.

**Supplementary Table 1.**
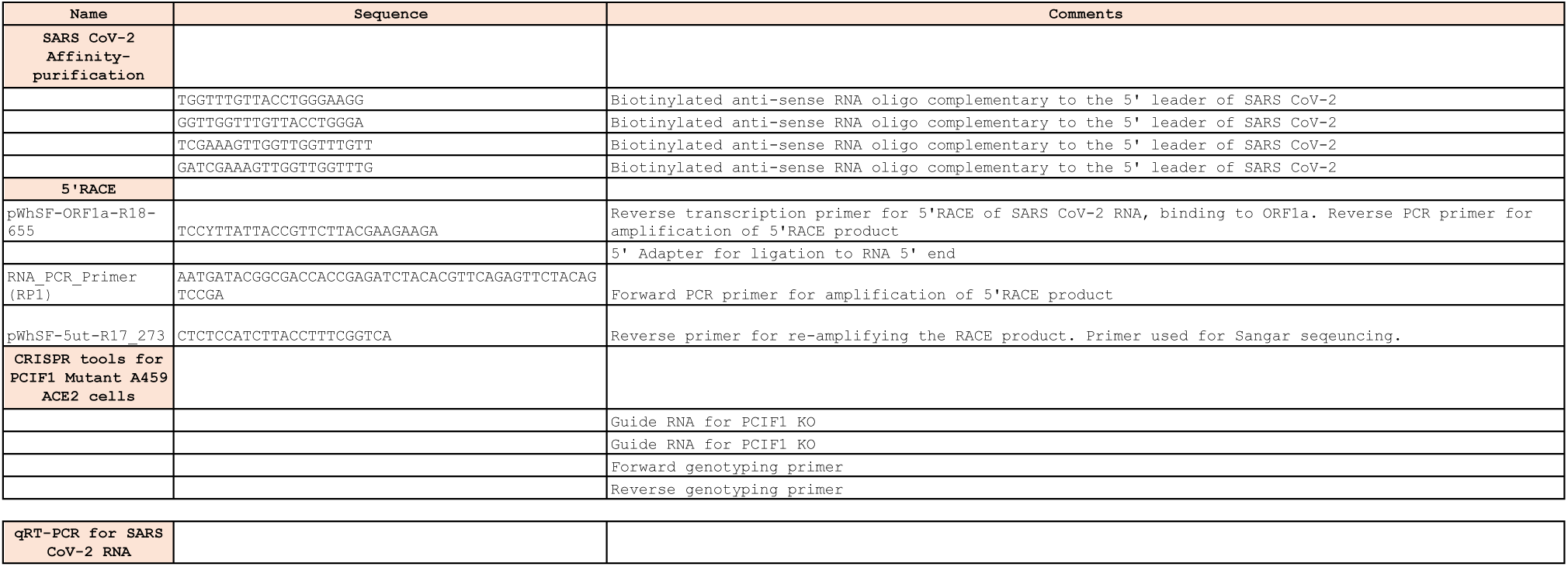
DNA and RNA oligonucleotides used in this study.

**Supplementary Table 2:**
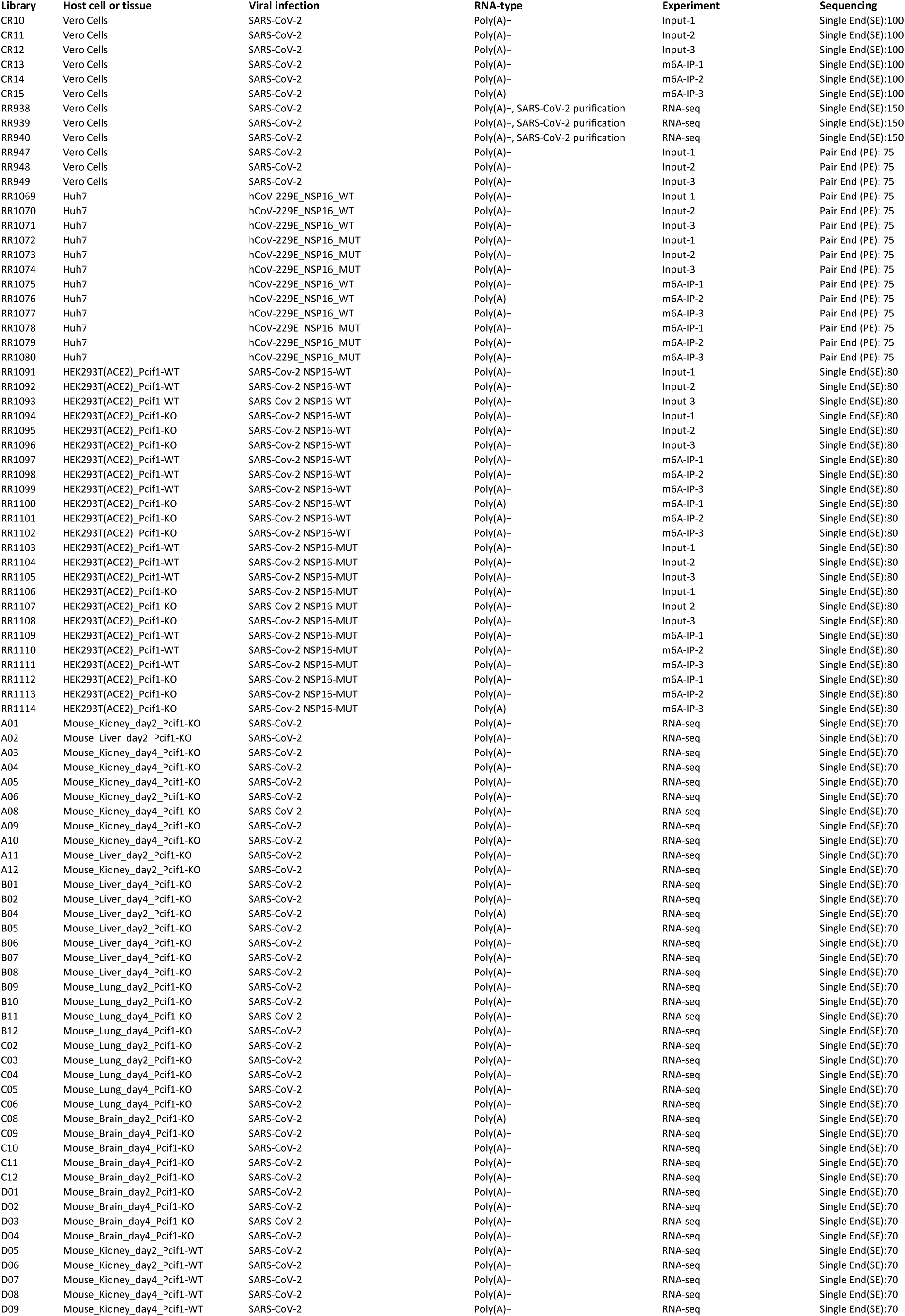
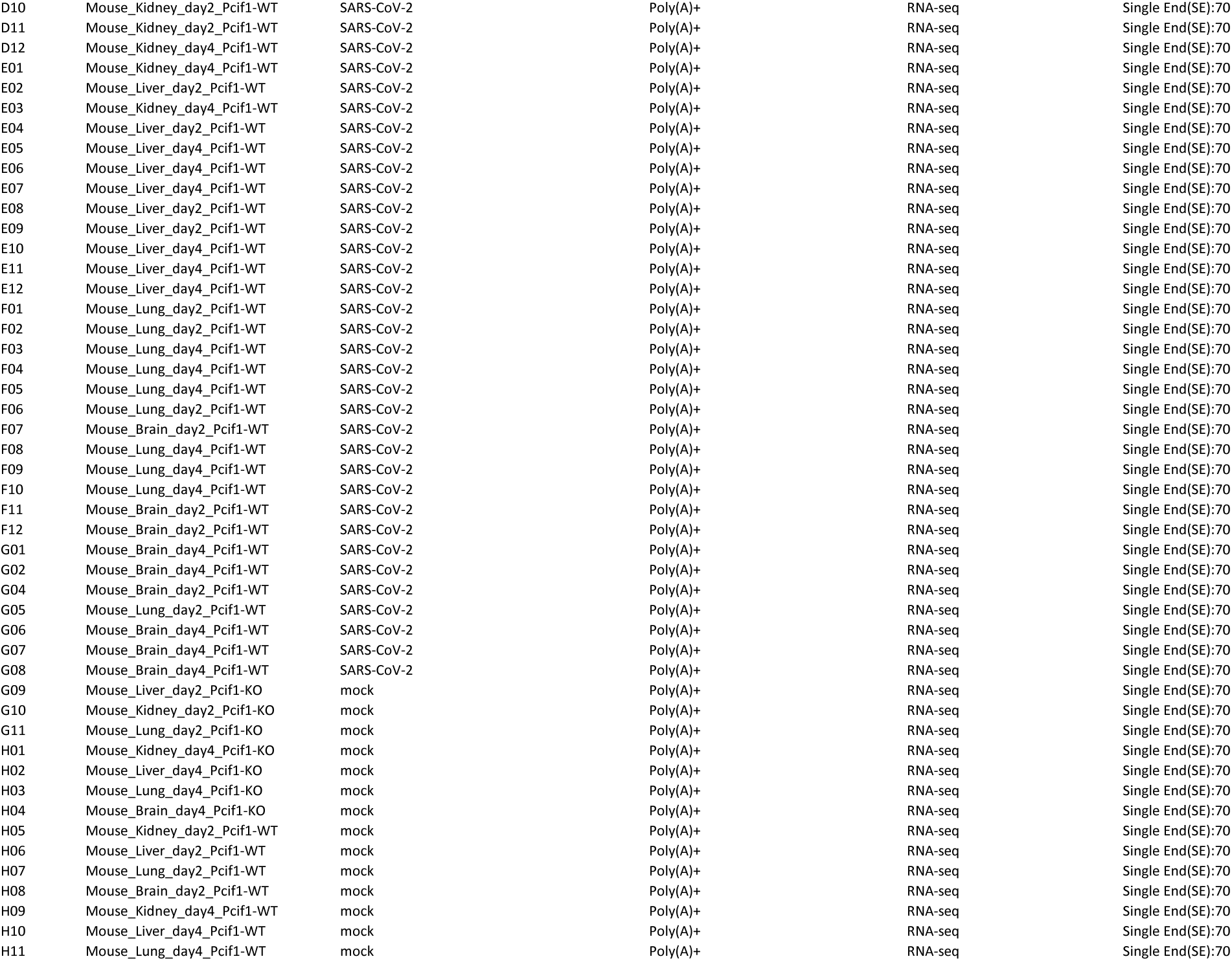
List of deep sequencing datasets generated in this study (GEO: GSEXXXX)

## References

1 Muthukrishnan, S., Moss, B., Cooper, J. & Maxwell, E. Influence of 5’-terminal cap structure on the initiation of translation of vaccinia virus mRNA. Journal of Biological Chemistry 253, 1710–1715 (1978).

2 Devarkar, S. C. et al. Structural basis for m7G recognition and 2′-O-methyl discrimination in capped RNAs by the innate immune receptor RIG-I. Proceedings of the National Academy of Sciences 113, 596–601 (2016).

3 Schuberth-Wagner, C. et al. A conserved histidine in the RNA sensor RIG-I controls immune tolerance to N1-2′ O-methylated self RNA. Immunity 43, 41–51 (2015).

4 Kowalinski, E. et al. Structural basis for the activation of innate immune pattern-recognition receptor RIG-I by viral RNA. Cell 147, 423–435 (2011).

5 Zust, R. et al. Ribose 2’-O-methylation provides a molecular signature for the distinction of self and non-self mRNA dependent on the RNA sensor Mda5. Nat Immunol 12, 137–143, doi:10.1038/ni.1979 (2011).

6 Akichika, S. et al. Cap-specific terminal N (6)-methylation of RNA by an RNA polymerase II-associated methyltransferase. Science 363, doi:10.1126/science.aav0080 (2019).

7 Sendinc, E. et al. PCIF1 Catalyzes m6Am mRNA Methylation to Regulate Gene Expression. Mol Cell 75, 620–630 e629, doi:10.1016/j.molcel.2019.05.030 (2019).

8 Boulias, K. et al. Identification of the m(6)Am Methyltransferase PCIF1 Reveals the Location and Functions of m(6)Am in the Transcriptome. Mol Cell 75, 631–643 e638, doi:10.1016/j.molcel.2019.06.006 (2019).

9 Pandey, R. R. et al. The Mammalian Cap-Specific m6Am RNA Methyltransferase PCIF1 Regulates Transcript Levels in Mouse Tissues. Cell reports 32, 108038 (2020).

10 V’kovski, P., Kratzel, A., Steiner, S., Stalder, H. & Thiel, V. Coronavirus biology and replication: implications for SARS-CoV-2. Nature Reviews Microbiology 19, 155–170 (2021).

11 Belanger, F., Stepinski, J., Darzynkiewicz, E. & Pelletier, J. Characterization of hMTr1, a human Cap1 2’-O-ribose methyltransferase. J Biol Chem 285, 33037–33044, doi:10.1074/jbc.M110.155283 (2010).

12 Werner, M. et al. 2′-O-ribose methylation of cap2 in human: function and evolution in a horizontally mobile family. Nucleic acids research 39, 4756–4768 (2011).

13 Inesta-Vaquera, F. & Cowling, V. H. Regulation and function of CMTR1-dependent mRNA cap methylation. Wiley Interdiscip Rev RNA 8, doi:10.1002/wrna.1450 (2017).

14 Liu, J. et al. A METTL3-METTL14 complex mediates mammalian nuclear RNA N6-adenosine methylation. Nat Chem Biol 10, 93–95, doi:10.1038/nchembio.1432 (2014).

15 Geula, S. et al. Stem cells. m6A mRNA methylation facilitates resolution of naive pluripotency toward differentiation. Science 347, 1002–1006, doi:10.1126/science.1261417 (2015).

16 Batista, P. J. et al. m(6)A RNA modification controls cell fate transition in mammalian embryonic stem cells. Cell Stem Cell 15, 707–719, doi:10.1016/j.stem.2014.09.019 (2014).

17 Schwartz, S. Cracking the epitranscriptome. RNA 22, 169–174 (2016).

18 Roignant, J. Y. & Soller, M. m(6)A in mRNA: An Ancient Mechanism for Fine-Tuning Gene Expression. Trends Genet 33, 380–390, doi:10.1016/j.tig.2017.04.003 (2017).

19 Wang, X. et al. N6-methyladenosine-dependent regulation of messenger RNA stability. Nature 505, 117–120, doi:10.1038/nature12730 (2014).

20 Mendel, M. et al. Splice site m6A methylation prevents binding of U2AF35 to inhibit RNA splicing. Cell (2021).

21 Zaccara, S. & Jaffrey, S. R. A unified model for the function of YTHDF proteins in regulating m6A-modified mRNA. Cell 181, 1582–1595. e1518 (2020).

22 Lasman, L. et al. Context-dependent functional compensation between Ythdf m6A reader proteins. Genes & development 34, 1373–1391 (2020).

23 Xiao, W. et al. Nuclear m(6)A Reader YTHDC1 Regulates mRNA Splicing. Mol Cell 61, 507–519, doi:10.1016/j.molcel.2016.01.012 (2016).

24 Daffis, S. et al. 2’-O methylation of the viral mRNA cap evades host restriction by IFIT family members. Nature 468, 452–456, doi:10.1038/nature09489 (2010).

25 Habjan, M. et al. Sequestration by IFIT1 impairs translation of 2′ O-unmethylated capped RNA. PLoS pathogens 9 (2013).

26 Lu, M. et al. N 6-methyladenosine modification enables viral RNA to escape recognition by RNA sensor RIG-I. Nature microbiology 5, 584–598 (2020).

27 Qiu, W. et al. N 6-methyladenosine RNA modification suppresses antiviral innate sensing pathways via reshaping double-stranded RNA. Nature communications 12, 1–16 (2021).

28 Zhang, X. et al. Methyltransferase-like 3 Modulates Severe Acute Respiratory Syndrome Coronavirus-2 RNA N6-Methyladenosine Modification and Replication. Mbio 12, e01067–01021 (2021).

29 Burgess, H. M. et al. Targeting the m6A RNA modification pathway blocks SARS-CoV-2 and HCoV-OC43 replication. Genes & development 35, 1005–1019 (2021).

30 Liu, J. e., et al. The m6A methylome of SARS-CoV-2 in host cells. Cell research 31, 404–414 (2021).

31 Li, N. et al. METTL3 regulates viral m6A RNA modification and host cell innate immune responses during SARS-CoV-2 infection. Cell reports 35, 109091 (2021).

32 Thi Nhu Thao, T., et al. Rapid reconstruction of SARS-CoV-2 using a synthetic genomics platform. Nature 582, 561–565, doi:10.1038/s41586-020-2294-9 (2020).

33 Chen, Y. et al. Biochemical and structural insights into the mechanisms of SARS coronavirus RNA ribose 2′-O-methylation by nsp16/nsp10 protein complex. PLoS pathogens 7 (2011).

34 Decroly, E. et al. Coronavirus nonstructural protein 16 is a cap-0 binding enzyme possessing (nucleoside-2’O)-methyltransferase activity. J Virol 82, 8071–8084, doi:10.1128/JVI.00407-08 (2008).

35 Hoffmann, M. et al. SARS-CoV-2 Cell Entry Depends on ACE2 and TMPRSS2 and Is Blocked by a Clinically Proven Protease Inhibitor. Cell, doi:10.1016/j.cell.2020.02.052 (2020).

36 Yount, B., Denison, M. R., Weiss, S. R. & Baric, R. S. Systematic assembly of a full-length infectious cDNA of mouse hepatitis virus strain A59. Journal of virology 76, 11065–11078 (2002).

37 Benoni, R. et al. Substrate specificity of SARS-CoV-2 nsp10-nsp16 methyltransferase. Viruses 13, 1722 (2021).

38 Leist, S. R. et al. A mouse-adapted SARS-CoV-2 induces acute lung injury and mortality in standard laboratory mice. Cell 183, 1070–1085. e1012 (2020).

39 Ulrich, L. et al. Enhanced fitness of SARS-CoV-2 variant of concern Alpha but not Beta. Nature, 1–10 (2021).

40 Winkler, E. S. et al. SARS-CoV-2 infection of human ACE2-transgenic mice causes severe lung inflammation and impaired function. Nature immunology 21, 1327–1335 (2020).

41 Park, G. J. et al. The mechanism of RNA capping by SARS-CoV-2. Nature, doi:10.1038/s41586-022-05185-z (2022).

42 Ivanov, K. A. et al. Multiple enzymatic activities associated with severe acute respiratory syndrome coronavirus helicase. Journal of virology 78, 5619–5632 (2004).

43 Walker, A. P. et al. The SARS-CoV-2 RNA polymerase is a viral RNA capping enzyme. Nucleic acids research 49, 13019–13030 (2021).

44 Yan, L. et al. Cryo-EM structure of an extended SARS-CoV-2 replication and transcription complex reveals an intermediate state in cap synthesis. Cell 184, 184–193. e110 (2021).

45 Chen, Y. et al. Functional screen reveals SARS coronavirus nonstructural protein nsp14 as a novel cap N7 methyltransferase. Proceedings of the National Academy of Sciences 106, 3484–3489 (2009).

46 Tartell, M. A. et al. Methylation of viral mRNA cap structures by PCIF1 attenuates the antiviral activity of interferon-β. Proceedings of the National Academy of Sciences 118 (2021).

47 Thao, T. T. N. et al. Rapid reconstruction of SARS-CoV-2 using a synthetic genomics platform. Nature 582, 561–565 (2020).

